# GCIB-SEM: A path to 10 nm isotropic imaging of cubic millimeter volumes

**DOI:** 10.1101/563239

**Authors:** K.J. Hayworth, D. Peale, M. Januszewski, G.W. Knott, Z. Lu, C.S. Xu, H.F. Hess

## Abstract

Focused Ion Beam Scanning Electron Microscopy (FIB-SEM) generates 3D datasets optimally suited for segmentation of cell ultrastructure and automated connectome tracing but is limited to small fields of view and is therefore incompatible with the new generation of ultrafast multibeam SEMs. In contrast, section-based techniques are multibeam-compatible but are limited in z-resolution making automatic segmentation of cellular ultrastructure difficult. Here we demonstrate a novel 3D electron microscopy technique, Gas Cluster Ion Beam SEM (GCIB-SEM), in which top-down, wide-area ion milling is performed on a series of thick sections, acquiring < 10 nm isotropic datasets of each which are then stitched together to span the full sectioned volume. Based on our results, incorporating GCIB-SEM into existing single beam and multibeam SEM workflows should be straightforward and should dramatically increase reliability while simultaneously improving z-resolution by a factor of 3 or more.

---

A wide range of biological research depends on visualizing the 3D ultrastructure of tissues with ^~^10 nm isotropic voxels. Although a variety of electron microscopic techniques can achieve this, none are well suited to image tissue volumes in the 1 mm^3^ range and above. Such large-volume ultrastructural imaging would be useful in the study of a variety of tissue systems, and it could truly revolutionize neuroscience allowing researchers to routinely image whole invertebrate nervous systems and substantial regions of small vertebrate brains. Perhaps most exciting is the prospect of directly testing long-standing theories which suggest learning and memory are encoded in changes to the structural connectome^1–6^, and which predict that simple memories might, in principle, be decoded from a map of connectivity and synaptic ultrastructure alone. This goal has been a key driver of innovation in the field^7^, but so far it has remained out of reach because of the requirement of automatically tracing volumes spanning many cubic millimeters. Diamond knife section-based techniques, i.e. ssTEM^8^, ATUM-SEM^9^, and S3EM^10^, theoretically have the imaging throughput needed because they can take advantage of fast, wide-area imaging technologies such as multibeam SEMs, especially the recently introduced 91 beam MultiSEM^11^, but they face the obstacle of how to consistently cut and collect tens of thousands of artifact-free sections thinly enough to provide the z-resolution needed for reliable automated tracing. Blockface techniques like SBEM^12^ and FIB-SEM^13^ already have the requisite z-resolution for automated tracing^14^, but so far they seem incompatible with high-throughput MultiSEM imaging. Here we demonstrate a new atomic cluster milling and imaging technique, Gas Cluster Ion Beam Scanning Electron Microscopy (GCIB-SEM), which represents a *hybrid* of FIB-SEM and diamond knife section-based techniques. In GCIB-SEM, fewer, much thicker sections are collected to span a given volume, then these thick sections are themselves ion milled in < 10 nm steps and volume imaged. This greatly improves sectioning reliability while simultaneously *improving* z-resolution and maintaining compatibility with MultiSEM imaging.

In FIB-SEM a tightly focused (^~^1 μm) beam of high-energy (30 kV) gallium ions is directed at an almost parallel angle (< 1°) to the surface of a tissue block, ablating material while avoiding deeper damage. Repeated FIB/SEM cycles produce a 3D stack, but milling artifacts accumulate in the direction of the FIB beam due to the shallow milling angle used, and these limit the area which can be imaged^13^. We reasoned that replacing FIB with a more ‘top-down’ approach might overcome this, but the energy of the ions would have to be lowered considerably. We therefore decided to try GCIB milling, a low-energy surface polishing method used in semiconductor fabrication and mass spectroscopy^15^. In GCIB, clusters of argon atoms are ionized and accelerated while a magnetic field selects clusters containing a particular number of atoms. In this way the average energy per atom can be tuned to just a few electron volts. Molecular dynamics simulations^16^ show that such clusters ‘splat’ against the surface, scattering atoms at glancing angles thereby removing asperities.

Our prototype system consists of a GCIB gun (GCIB-10s from Ionoptika) mounted to a single beam SEM (Ultra SEM from Zeiss) (**Supplementary Fig. 1**). We first tested milling a variety of ultrathin tissue sections, exploring a range of angles, cluster sizes and energies, and embedding resins. We SEM imaged milled surfaces with 1.2 kV landing energy (chosen to provide adequate z-resolution) using energy selective backscatter (ESB) and in-lens secondary electron (InLens-SE) detectors. ESB (500 V filtering, see **Methods**) produced high quality images under virtually all milling conditions, but InLens-SE showed rough surfaces in some. InLens-SE detection is much more sensitive to surface topography and charging than ESB detection but approximates MultiSEM imaging^11^. Epon samples showed rough InLens images under all but the shallowest milling angles. Durcupan and Sprurr’s samples gave high-quality InLens-SE images at glancing angles up to 30° (**Supplementary Fig. 2**).

Sections thicker than 100 nm charged during imaging, and GCIB milling did not correct this (unlike FIB-SEM). Charging lessened with extended imaging and we reasoned that this was due to the slow conversion of the polymer embedding into a more conductive amorphous carbon^17^. We subsequently demonstrated that such electron irradiation could be used to make sections at least 10 μm thick sufficiently conductive to allow quality InLens-SE imaging (**Supplementary Fig. 3,4**). We found that a dose of ^~^0.5×10^27^ eV/cm^3^ was needed to completely eliminate charging. GCIB milling of irradiated sections remained sufficiently smooth to produce 3D volume images but the sputter removal rate of irradiated regions dropped by up to a factor of six (**Supplementary Fig. 5**). Our current ‘optimized’ protocol recommends irradiating to a dose of 0.5×10^27^ eV/cm^3^, then milling with 10 kV Ar2000 clusters at a glancing angle of 30° while the sample is rotating. **Fig. 1a** outlines this process.

**Figure 1.**
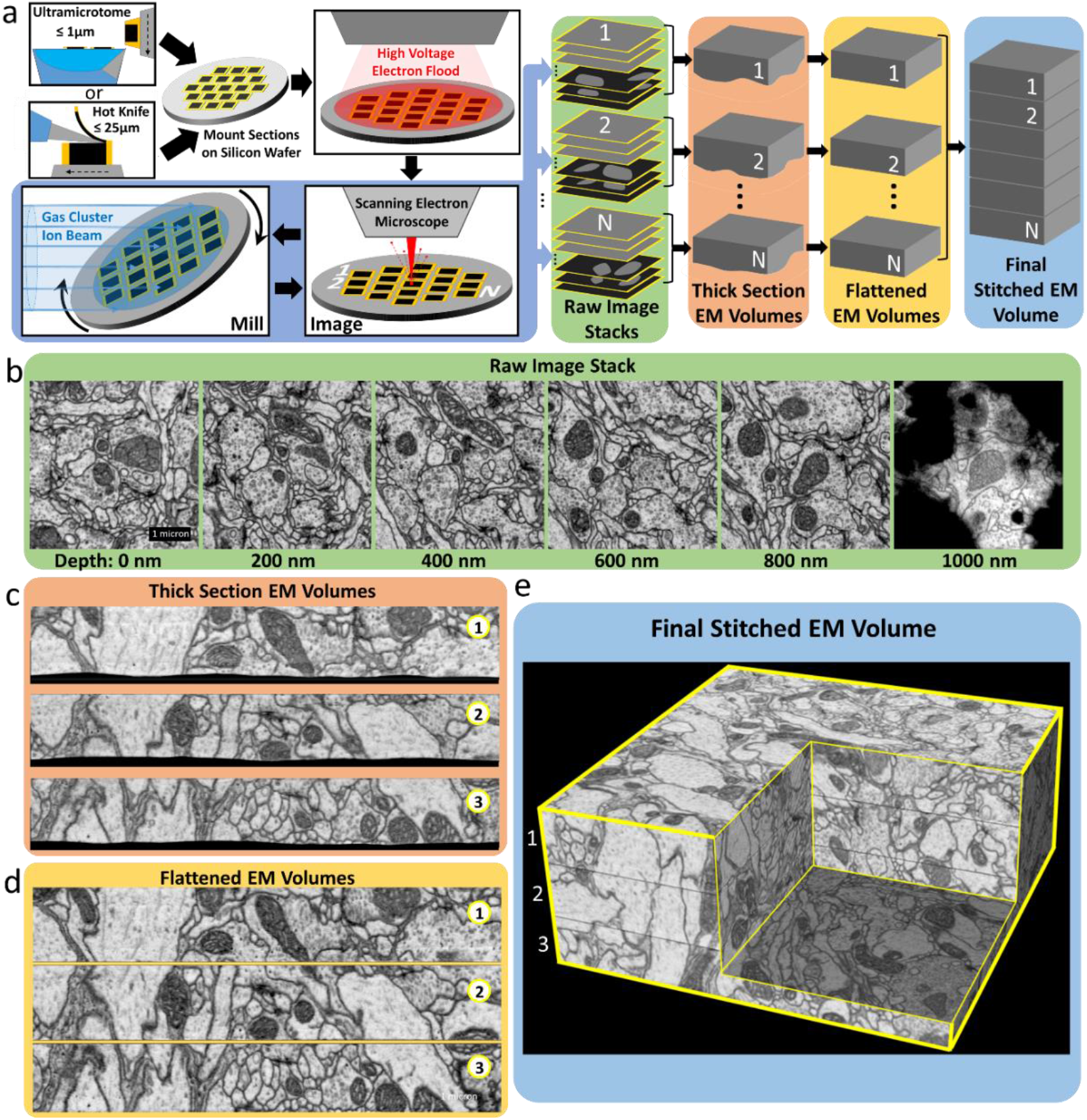
GCIB-SEM overview. **(a)** Pictorial diagram of the GCIB-SEM process. A series of ‘thick’ sections (≤ 1 μm) collected by traditional ultramicrotomy, or a series of ‘ultrathick’ sections (≤ 25 μm) collected by ‘hot knife’ sectioning, are mounted on a silicon wafer and then irradiated with a high-voltage electron flood gun to increase their conductivity. The wafer is then shuttled repeatedly between SEM-imaging and GCIB-milling steps creating a set of raw image stacks, each spanning the depth of one of the thick sections. The raw images are first aligned to create 3D volumes of each individual thick section, then these volumes are “flattened” and stitched together to create a single 3D volume spanning the entire sequence of thick sections. Numbers 1,2,…N denote individual thick sections. **(b-e)** Example images from a GCIB-SEM run performed on three sequential 1 μm thick sections of Durcupan-embedded *Drosophila* brain. All images acquired using InLens-SE detection. **(b)** Selected raw images, one every 200 nm of milled depth, spanning a single thick section. The last image shows milling breaking through to the gold substrate (black pixels). **(c)** Z-reslice through the image stacks of all three thick sections prior to flattening and stitching. Black pixels at bottom are gold substrate. **(d)** Z-reslice of same stack after flattening. These are shown aligned to demonstrate how the flattening procedure allows for tracing of processes across thick section boundaries. **(e)** Stitched stack shown as 3D volume. Numbers 1,2,3 in **c-e** denote individual thick sections.

To test, we sectioned a Durcupan-embedded *Drosophila* brain and collected three sequential 1 μm sections onto gold-coated silicon. We performed ^~^250 GCIB/SEM cycles (using InLens-SE detection) completely milling through all three sections (6×6×4 nm voxels). **Fig. 1b** shows images of one section at 200 nm intervals. **Fig. 1c** shows a Z-reslice through the image stacks of all three sections showing an unevenness at the bottoms of each section, the result of milling rate variance (^~^10%). We choose to collect on gold-coated silicon because the InLens-SE and ESB signals for gold are beyond the maximum signals produced by heavy-metal stain. Remarkably this 10% milling variance is sufficiently small and sufficiently spatially uniform that these variances could be corrected by software. We wrote algorithms to find and ‘flatten’ this unevenness (see **Methods**). Once flattened (**Fig. 1d**), we stitched the sections into a single volume (**Fig. 1e**). Membranes and synapses are clear and resemble those we typically see using FIB-SEM. A close inspection (**Supplementary Video 1**) shows some slight artifacts relative to FIB-SEM, particularly a light texture of milling streaks is visible in the Z-reslice.

This sample was embedded in Durcupan resin which appears to be insufficiently resilient to be sectioned this thick (diamond cut surface showed some damage), so we have now switched to Spurr’s embedding which can be room-temperature sectioned with high quality surfaces at up to 1 μm, and can be hot knife sectioned at up to 25 μm. **Fig. 2a** shows a Z-reslice of a GCIB-SEM volume (using InLens-SE detection) covering three sequential 500 nm thick sections of Spurr’s-embedded mouse cortex (8×8×6 nm voxels). Ultrastructural features like post synaptic densities and vesicles are clearly defined. **Fig. 2b** shows the simultaneously-acquired ESB volume which looks almost identical under these milling conditions. Cut surfaces were smooth and were easily flattened and stitched with an estimated loss between sections of ^~^30 nm (see **Methods**), and visual tracing of all neuronal processes appeared straightforward (**Fig. 2c**, **Supplementary Video 2**). These datasets demonstrate that wide-area GCIB milling and InLens-SE detection can be performed over multiple sequential thick sections and can produce datasets approaching the quality of FIB-SEM.

**Figure 2:**
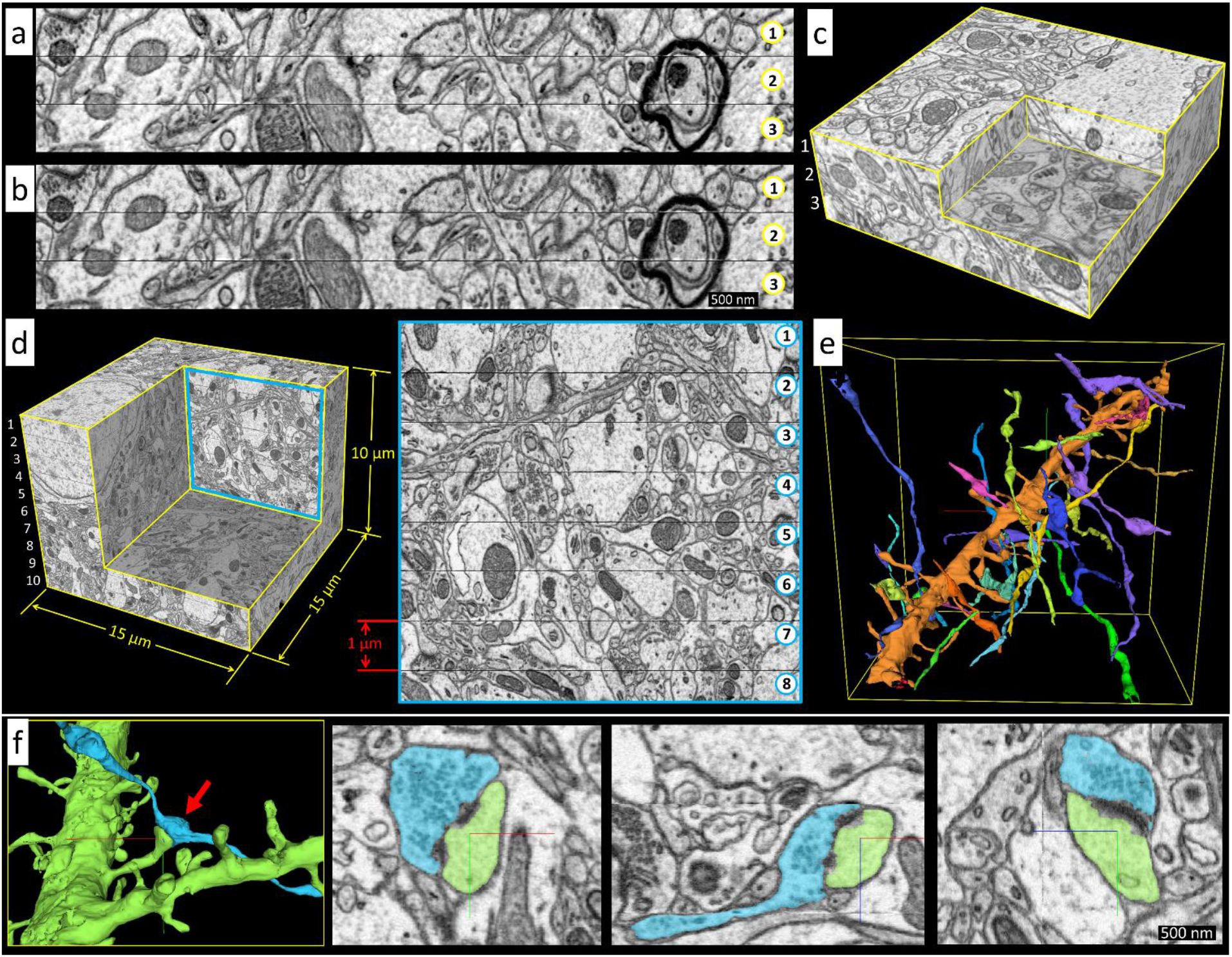
Examples of GCIB-SEM imaging. **(a-c)** Images from a GCIB-SEM run performed on three sequential 500nm thick sections of mouse cortex. **(a)** Z-reslice through the InLens-SE-detected stack. **(b)** Z-reslice through the ESB-detected stack. Note that InLens-SE and ESB stacks are similar under the milling conditions used. **(c)** InLens-SE stack shown as 3D volume. Numbers denote individual thick sections. **(d)** GCIB-SEM 3D volume spanning ten sequential 1 μm sections of mouse cortex, along with example cross section (blue framed region, numbers denote individual thick sections, ESB detection shown). **(e)** Tracing of a single spiny dendritic process (orange) spanning the volume along with axons synapsing on it. **(f)** XY, XZ, and YZ views through a single spiny synapse (red arrow) from this volume.

To address the applicability of GCIB-SEM to mammalian connectomics we sectioned Spurr’s-embedded mouse cortex at 1 μm and collected ten sequential sections, then performed ^~^170 GCIB/SEM cycles (8×8×6 nm voxels) completely milling through all ten sections. We computationally flattened each section and stitched them together into the single volume shown in **Fig. 2d**, **Supplementary Fig. 6–11**, and **Supplementary Video 3**. The cut surfaces of these Spurr’s-embedded sections were of high quality and we estimated the loss between sections at ^~^30 nm. A test segmentation of the ESB stack was performed using a flood-filling neural network^14^ that had previously been trained on a quite different SBEM dataset and then ‘bootstrap-trained’ on this dataset (see **Methods**). **Fig. 2e** shows an example spiny dendrite spanning the ten section GCIB-SEM volume along with synapsing axons. **Fig. 2f** shows views through a single spiny synapse.

Cutting at 1 μm appears to be approaching the limit for room temperature sectioning of Spurr’s since we saw significant surface damage when we attempted sectioning at 2 μm. Cutting much thinner (e.g. 150 nm) resulted in less loss and higher quality surfaces (see **Methods**), which may be preferred despite the fact that it results in more stitch boundaries. An alternative is hot knife ultrathick sectioning which uses a heated diamond knife and oil lubrication to minimize stresses on the tissue^18^.

To test this, we ‘hot knife’ sectioned Spurr’s-embedded mouse cortex at 10 μm thickness and flat embedded two sequential sections against gold-coated silicon. The resulting GCIB-SEM volume (10×10×12 nm voxels) spanning the two hot knife sections is shown in **Fig. 3a** and a test segmentation based on the InLens-SE stack using a flood-filling network is shown in **Fig. 3b**. **Supplementary Figs. 12–16** show Z-reslice views of the GCIB-SEM stack spanning the two 10 μm thick sections before and after flattening and stitching as well as additional segmentation examples. The cut surfaces were of high quality but the flat embedding procedure resulted in some contamination. We estimated the loss between sections at ^~^30 nm (see **Methods**). **Supplementary Video 4** shows a side-by-side comparison of the ESB vs. InLens-SE volumes (also **Fig. 2c,d**).

**Figure 3:**
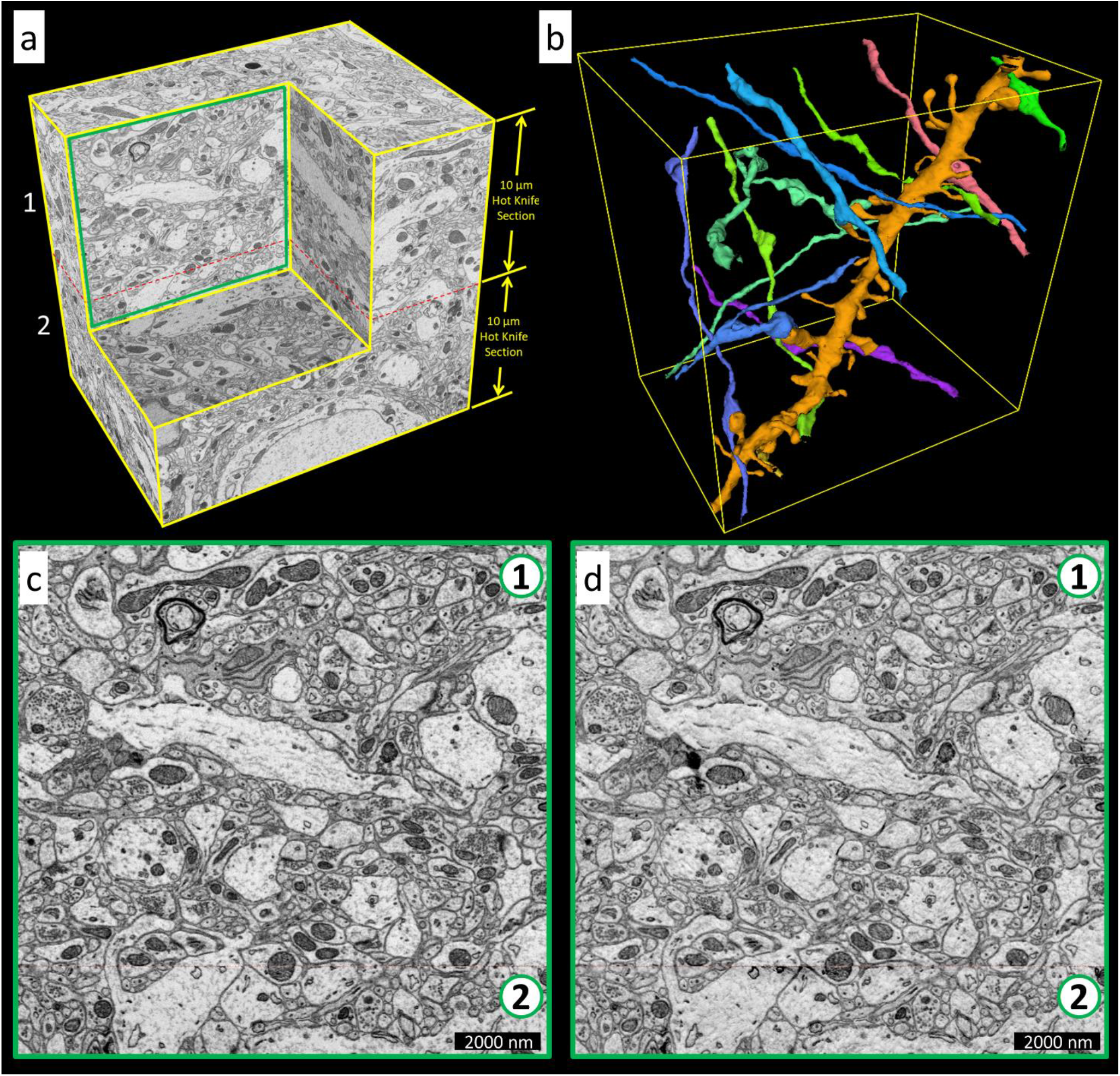
GCIB-SEM imaging of two sequential 10 μm “ultrathick” hot knife cut sections. **(a)** GCIB-SEM volume (ESB detection shown). **(b)** Tracing of a single spiny dendritic process (orange) spanning the volume along with axons synapsing on it. This tracing is based on a segmentation of the InLens-SE detected volume. **(c)** Example z-reslice (green framed region in **a)** showing the top section’s full 10 μm depth and its stitch plane (red dashed line) to the bottom section. **(d)** Same z-reslice but from the InLens-SE detected volume.

We noticed that GCIB milling produces a slight drop in the InLens-SE membrane contrast which progresses over the first ^~^500 nm of milling. We speculate this is the result of a few nm thick layer of disrupted surface material persisting in equilibrium with milling which changes SE yield. Initial electron irradiation also produces a slight drop in InLens-SE contrast. These effects are quantified in **Supplementary Fig. 17**. We also verified that GCIB-SEM imaging was generally compatible with ATUM tape collection^9^ by imaging two 1 μm thick sections collected on copper coated tape (**Supplementary Fig. 18**).

Finally, we verified that the InLens-SE detection used in our prototype was an adequate proxy for MultiSEM imaging. We GCIB-SEM milled and imaged the first 250 nm of a 500 nm section of mouse cortex then shipped the half-milled section to Zeiss for imaging on their MultiSEM system. MultiSEM images of the GCIB-milled surface (acquired with similar electron dose) were essentially identical to our InLens-SE images of the same surface (**Supplementary Figs. 19,20**).

We see Serial Thick-Section GCIB-SEM as a potential route to ‘industrial-scale’ 3D EM and connectomics as it addresses key limitations of existing techniques: GCIB-SEM resolution is not limited by section thickness, and imaging dose is not limited by interactions with sectioning^19^, so resolution and dose can be adjusted as needed to achieve fully automated segmentation. GCIB-SEM utilizes thick sectioning which should be reliable even over arbitrarily large volumes. GCIB is dramatically less sensitive to beam position and focus than FIB, allowing lossless restarts. The gas cluster source, unlike FIB, does not require periodic reheating and is comparatively simple and reliable. GCIB-milling is wide-area and fast (up to 450 μm^3^/s, 1 mm^3^/month, see **Methods**), and, if necessary, can be performed across multiple GCIBs in parallel with imaging when sections are spread across multiple wafers. This should allow close to 100% utilization of MultiSEM imaging time and should allow projected^20^ MultiSEM improvements to be taken full advantage of. We envision a day when acquiring automatically segmented EM volumes of large tissue samples (e.g. whole invertebrate and small vertebrate brains) is routine; where such volumes are reliably thick sectioned and spread across multiple wafers which are then robotically shuttled between multiple GCIB-milling and MultiSEM-imaging stations in a setting reminiscent of today’s semiconductor fabrication plants.

## Acknowledgments

We thank Jorgen Kornfeld for allowing the use of a flood-filling network which was trained on one of his SBEM datasets. We thank Anna Eberle for MultiSEM imaging our GCIB-SEM samples. We thank Yoshiyuki Kubota for providing the copper tape used in our ATUM collection tests. We thank Winfried Denk and Maria Kormacheva for useful discussions.

## Online Methods

### Sample preparation

All experimental protocols were conducted according to US National Institutes of Health guidelines for animal research and were approved by the Institutional Animal Care and Use Committee at Janelia Research Campus. A three-month-old adult C57/bl6 mouse was euthanized with an overdose of isoflurane and immediately perfused via the heart with a buffered solution of 1.25 % glutaraldehyde and 4 % paraformaldehyde (0.1M PB pH 7.4). Two hours after perfusion the brain was removed and left in the same fix overnight at 4°C. Coronal sections of the brain were then cut with a vibratome (Leica VT1200). These were stained with 1.5% potassium ferrocyanide, on ice, followed by 2% osmium, each diluted in phosphate buffer. Sections were stained in 1% thiocarbohydrazide for 20 minutes before transferring them into 2% osmium tetroxide (Agar Scientific) for a further 30 minutes. They were then placed in 1% uranyl acetate at 4 °C overnight then in lead aspartate solution at 50 °C for 2 hours. The sections were finally dehydrated in increasing concentrations of alcohol, for 5 minutes each change and then transferred to increasing concentrations of epoxy resin (Spurs, EMS) until 100%. These were placed between two glass slides, coated with mold separating agent (Glorex, Switzerland), and hardened at 60 ° C for 24 h.

A six-day-old male adult Canton S G1 x w1118 fly of the *Drosophila* was GCIB-SEM imaged producing the micrographs shown in **Fig. 1b-e**. The preparation method used was a heavy metal contrast enhancement on C-PLT, an optimized preparation protocol for *Drosophila* brains with optimal contrast and morphological preservation for FIB-SEM. Chemical/ Progressive Lowering of Temperature (C-PLT) fixation/dehydration with *en bloc* staining is a modified conventional chemical fixation method^13,18^. Isolated whole brains were fixed in 2.5% formaldehyde and 2.5% glutaraldehyde in 0.1 M phosphate buffer at pH 7.4 for 2 hours at 22 °C. After washing, the tissues were post-fixed in 0.5% osmium tetroxide in 0.05M sodium cacodylate buffer for 40 min and then treated with 0.8% potassium ferricyanide in buffer for 2 hours at 4 °C. After washing, tissue was incubated with 0.5% aqueous uranyl acetate for 30 min at 4 °C then followed by lead aspartate *en bloc* staining at 4 °C for overnight. A PLT procedure started from 1 °C when the tissues were transferred into 10% ethanol after secondary osmication with 0.8% osmium tetroxide for 20 min at 4 °C. The temperature was progressively decreased to −25 °C while the ethanol concentration was gradually increased to 97%. The tissue was incubated in 1% osmium tetroxide and 0.2% uranyl acetate in ethanol for 32 hours at −25 °C. After PLT and low temperature incubation, the temperature was increased to 22 °C, and tissues were rinsed in pure ethanol following by acetone, then infiltrated and embedded in Durcupan (ACM Fluka).

### Sectioning, Irradiation, Milling and Imaging

#### *Drosophila* brain (Fig. 1b-e)

We sectioned a Durcupan-embedded *Drosophila* brain at 1 μm thickness on an ultramicrotome (Leica UC7) using a room temperature diamond knife (Diatome, 35° clearance angle) and collected three sequential sections from the water boat onto a gold-coated silicon wafer. Silicon wafers used for this and other runs were pre-diced 5 x 7 mm p-type silicon support chips (Ted Pella). We deposited 5 nm of chromium (to promote gold adhesion) followed by 50 nm of gold onto the silicon surface using a Precision Etching and Coating System (Gatan). We then glow-discharged the gold surface just prior to collection to promote wetting. Electron irradiation parameters used: 10,000 μm^2^ area, 2 hrs at 6 kV, 4 hrs at 10 kV, 1×10^27^ eV/cm^3^ total dose. Electron irradiation for this and other samples was performed using the SEM’s beam set to its highest aperture, defocused and scanned across the area to be irradiated. GCIB parameters used: 22nA of Ar2000 at 10 kV spread over a 10 mm^2^ area, 21° glancing angle, 360° rotation (3 steps), 900 s per mill cycle giving 4 nm of removal per image (44 μm^3^/s milling rate). GCIB milling for this and other samples was performed by slightly defocusing the GCIB beam (^~^0.5 mm) and scanning the area to be milled with a 50 x 50 position grid. SEM parameters used: 1.2 kV, 2 nA electron beam, InLens-SE detection, 6 nm pixels, 2 MHz acquisition rate.

#### Mouse cortex (Fig. 2a-c)

We sectioned Spurr’s embedded mouse cortex at 500 nm thickness on an ultramicrotome (Leica UC7) using a room temperature diamond knife (Diatome, 35° clearance angle) and collected three sequential sections from the water boat onto a gold-coated silicon wafer. Electron irradiation parameters used: 10,000 μm^2^ area, 4 hrs at 6 kV, 0.8×10^27^ eV/cm^3^ total dose. GCIB parameters used: 22 nA of Ar2000 at 10 kV spread over a 9.7 mm^2^ area, 30° glancing angle, 360° rotation (3 steps), 240 s per mill cycle giving 6 nm of removal per image (240 μm^3^/s milling rate). SEM parameters used: 1.2 kV, 2 nA electron beam, simultaneous ESB (500 V filtering) and InLens-SE detection, 8 nm pixels, 1.25 MHz acquisition rate.

#### Mouse cortex (Fig 2d-f)

We sectioned Spurr’s embedded mouse cortex at 1 μm thickness on an ultramicrotome (Leica UC7) using a room temperature diamond knife (Diatome, 35° clearance angle) and collected ten sequential sections onto a gold-coated silicon wafer. Electron irradiation parameters used: 10,000 μm^2^ area, 1.6 hrs at 8kV, 2.4×10^26^ eV/cm^3^ total dose. GCIB parameters used: 22nA of Ar2000 at 10kV spread over a 18 mm^2^ area, 30° glancing angle, 360° rotation (3 steps), 240s per mill cycle giving 6nm of removal per image (450 μm^3^/s milling rate). SEM parameters used: 1.2kV, 2nA, simultaneous ESB (500 V filtering) and InLens-SE detection, 8nm pixels, 1.25MHz acquisition rate.

#### Mouse cortex ‘hot knife’ sectioned (Fig 3)

We ‘hot knife’ sectioned Spurr’s embedded mouse cortex at 10 μm thickness using the method described in (Hayworth et al. 2015)^18^. Hot knife parameters used: Diamond knife (Diatome cryo, 25° clearance angle) lubricated with 0.22 μm filtered tapping oil (Master Plumber), 65°C knife temperature, no oscillation, 0.1 mm/s cutting speed. Two sequential sections were collected, washed of oil by dipping repeatedly in uncured Durcupan resin, and flat embedded against a gold coated silicon wafer (while still covered in uncured Durcupan) by sandwiching sections between the wafer and a glass optical flat that had a Kapton release film taped over it. This sandwich was put in a 65°C oven to cure under a weight. This flat embedding procedure resulted in the two thick sections having a thin covering of Durcupan that had to be GCIB-milled to reveal their tissue surfaces (see text). A scalpel and laser ablation were used to remove large regions of the surrounding Durcupan ‘flash’ to prevent peripheral charging, then matching regions in each thick section were electron irradiated to make them conductive. Electron irradiation parameters used: 110,000 μm^2^ area, 117 hrs at 30 kV, 0.9×10^27^ eV/cm^3^ total dose. GCIB parameters used: 22nA of Ar2000 at 10kV spread over a 13 mm^2^ area, 30° glancing angle, 360° rotation (6 steps), 360 s per mill cycle giving 12 nm of removal per image (400 μm^3^/s milling rate). SEM parameters used: 1.2 kV, 2 nA electron beam, simultaneous ESB (500 V filtering) and InLens-SE detection, 10 nm pixels, 1.25 MHz acquisition rate.

#### Mouse cortex (MultiSEM test sample, Supplementary Figs. 19, 20)

We sectioned Spurr’s embedded mouse cortex at 500 nm thickness on an ultramicrotome (Leica UC7) using a room temperature diamond knife (Diatome, 35° clearance angle) and collected a single section from the water boat onto a gold-coated silicon wafer. A region was electron irradiated to make it conductive, then GCIB-SEM imaged to a depth of ^~^250 nm before shipping to Zeiss for MultiSEM imaging tests. Electron irradiation parameters used: 188,000 μm^2^ area, 89 hrs at 6 kV, 0.9×10^27^ eV/cm^3^ total dose. GCIB parameters used: 32 nA of Ar2000 at 10kV spread over a 8 mm^2^ area, 30° glancing angle, 360° rotation (3 steps), 360 s per mill cycle giving ^~^12 nm of removal per image. SEM parameters used for GCIB-SEM imaging on Janelia Ultra SEM: 1.2 kV, 2 nA electron beam, InLens-SE detection, 8 nm pixels, 1.25 MHz acquisition rate.

### Computational flattening and stitching of sections

Processing of all GCIB-SEM runs proceeded in three steps (overviewed in **Fig. 1**): 1.) Align the SEM images of each thick section individually to generate a set of 3D volumes, one for each thick section. 2.) Computationally flatten the bottoms of each section individually. 3.) Align and stitch these 3D volumes together to form a final spanning 3D volume. We used the SIFT and BUnwarpJ alignment tools in the Fiji software^21^ to perform steps #1 and #3. For step #2 we wrote a custom Matlab script to perform flattening.

To perform flattening we first identified the grayscale threshold that best separated tissue pixels from substrate pixels, which could be reliably distinguished by grayscale as long as the thick sections were collected on a gold substrate. Our flattening script used this grayscale threshold to determine, for each x,y pixel position, the image index (z-position) where the tissue broke through to the substrate, creating a “bottom index map”. This “bottom index map” was lightly smoothed spatially and then used to stretch (with linear interpolation) each column of pixels in the z-direction so that all substrate breakthroughs would occur in a single z-plane.

We have provided this custom Matlab flattening script in the **Supplementary Software** along with an example dataset and a brief manual describing its operation.

### Section-to-section loss estimation

We estimated section-to-section tissue losses for each of the GCIB-SEM runs described in the text as well as for a test run performed with a series of 150 nm thick sections. **Supplementary Figs. 21–25** summarize these results.

Our method for estimating section-to-section loss was as follows: First the z-thickness of each GCIB-SEM mill step was estimated by dividing the microtome-set cutting thickness by the number of mill steps spanning a thick section. Then each image in the final stitched GCIB-SEM stack covering multiple serial thick sections was first spatially filtered (using FIJI) with a 2D Gaussian (r = 4 pixels), and a matrix of mean pixel-wise image differences (absolute valued) spanning the full GCIB-SEM volume was computed using a custom Matlab script. We used this to plot the expected image difference vs. mill steps (shown in each supplementary figure). The images right next to the cut surfaces display differences in contrast and features relative to images in the interior of a stack. For example, the first few images of each thick section often have somewhat higher contrast which diminishes after milling (see **Supplementary Fig. 17**), and the last few images of a flattened stack show contrast and texture artifacts from the flattening algorithm where pixel grayscale values start to blend with the gold substrate. Looking at these ‘boundary’ images makes clear that they contain information crucial for tracing but directly computing the image difference between such ‘top’ and ‘bottom’ images would lead to an erroneously large estimated loss dominated by such contrast and flattening-artifact effects. Instead we picked images slightly removed from the boundary to compare and computed the loss based on these.

For example, the GCIB-SEM dataset shown in **Fig. 2a-c** spanned three 500 nm microtome cut sections. The flattened stack for the first thick section from this run contained 81 images giving an estimated mill step size of 500 nm / 81 = 6.2 nm. The computed image difference matrix is shown in **SOM Fig. 22** along with a plot of the expected image difference vs. mill steps derived from looking at only ‘interior’ images. The boundary between the second and third thick section occurred between image #162 and #163 in this stack, but image #162 was not sufficiently clean for direct comparison due to the contrast and flattening artifacts described above. Instead we backed up to image #159 (which was clean) and compared it to image #163. The difference between these images (according to the computed mean difference matrix) was 14.5 units which corresponded to 8.7 mill steps according to the plot. This implies that there is an equivalent of 8.7 − 3 = 5.7 mill steps between #162 and #163. But if there was no loss of tissue during sectioning then there should only be 1 mill step between #162 and #163. So the loss is equivalent to 5.7 − 1 = 4.7 mill steps or 4.7 * 6.2 nm = 29 nm.

As described, the flattening procedure results in boundary images containing artifacts and, depending circumstances (for example, how sensitive one’s segmentation algorithm is to noise), one may decide to discard one or more of the boundary images. To account for this **Supplementary Figs. 21–25** display both the boundary image we based our loss estimate on and the next one that would be used if that one was discarded along with loss estimates for both.

### Flood-filling network segmentation

As described in the text, test segmentations were performed on an ESB-detected GCIB-SEM volume spanning ten 1 μm sections of mouse cortex which was acquired with 8×8×6 nm voxels (**Fig. 2d-f**, **Supplementary Figs. 6–11**) and on an InLens-SE-detected GCIB-SEM volume spanning two 10 μm ‘hot knife’ sections of mouse cortex which was acquired with 10×10×12 nm voxels (**Fig. 3**, **Supplementary Figs. 12–16**). The two GCIB-SEM stacks were segmented with flood-filling networks (FFN)^14^. Instead of producing de novo dedicated volumetric training data for these GCIB-SEM volumes, an existing model trained on 10×10×25 nm SBEM data of zebra finch Area X tissue (provided by Jorgen Kornfeld) was used to bootstrap the segmentation, and object-based proofreading was utilized to correct errors as described next.

Using the ESB-detected GCIB-SEM volume spanning ten 1 μm sections, a prototype variant of the CycleGAN-based transfer procedure described in (Januszewski & Jain 2019)^22^ was first applied between (a) 20×20×25 nm SBEM and 16×16×24 nm GCIB-SEM data (downsampled from its native 8×8×6 nm acquisition resolution), as well as 10×10×25 nm SBEM and (b) 16×16×12 nm, and (c) 8×8×18 nm GCIB-SEM data. Reduced-resolution downsampled variants of the GCIB-SEM volume were obtained via area-averaged resampling of the full resolution data.

Segmentations obtained for multiple CycleGAN checkpoints from the (a) and (b) variants were screened for merge errors, and combined with oversegmentation consensus after upsampling to a common resolution. 12 neurite fragments were manually assembled from correct fragments in the base segmentation. These fragments were then used to train an FFN model for 16×16×12 nm GCIB-SEM data, with loss masking in 2 images before and after every 1 μm thick section seam. The segmentation from this new model correctly resolved the larger structures in the volume, but the model could not track finer processes due to insufficient downsample resolution of input data and a lack of representative training examples. To compensate for that, oversegmentation consensus between the FFN-generated reduced resolution segmentation, and the full resolution segmentation from SBEM-transfer (see (c) above) was computed, an additional 13 neurites were reconstructed by manual fragment assembly to form the training set for a full-resolution GCIB FFN model. Once trained, a segmentation was generated, 450 objects were proofread and selected for another round of training. All objects larger than 100k voxels were inspected in the final segmentation volume and split errors were manually corrected where necessary.

A similar procedure was applied to segment the second GCIB-SEM volume (InLens-SE-detected GCIB-SEM volume spanning two 10 μm ‘hot knife’ sections, acquired at 10×10×12 nm voxel resolution). First, the network trained on the first GCIB-SEM volume at 8×8×6 nm resolution was transferred to the InLens-SE 10×10×12 nm stack with the help of a CycleGAN. No attempt was made to correct the mismatched voxel resolution.

Candidate segmentations were generated for multiple checkpoints of the CycleGAN using forward-backward oversegmentation consensus^14^. The following iterative procedure was then applied to identify the best checkpoint. Starting with a random checkpoint, 3D meshes of objects in the segmentation were screened for mergers in descending order of object voxel count until 10 merge locations were found and added to a list of known mergers. The segmentations were then evaluated using all recorded merge locations, and the segmentation with the fewest mergers was screened again. Screening was terminated once a single segmentation without known errors remained, at which point 61 distinct merge errors were recorded. FFN agglomeration with standard settings^14^ was then applied to the selected segmentation, with all merge pairs from agglomeration subjected to manual review. All objects larger than 100k voxels after agglomeration and touching 0 or 1 faces of the volume were then manually inspected, and if necessary connected to other objects within the volume. For the final segmentation, an FFN model was retrained on this proofread segmentation with the voxelwise loss computed only over voxels belonging to objects larger than 100k voxels.

The FFN network architecture and training procedures described in (Januszewski et al. 2018)^14^ were used everywhere, but the field of view was extended to (33, 33, 33) voxels to account for the more isotropic resolution of the GCIB-SEM data, and the depth of the network for the second GCIB-SEM volume was extended to 12 residual modules. The manual inspections performed did not reveal any objects that could not be unambiguously traced. No manual voxel-level corrections were performed at any step of the reconstruction.

## SUPPLEMENTARY FIGURES

**SOM Figure 1.**
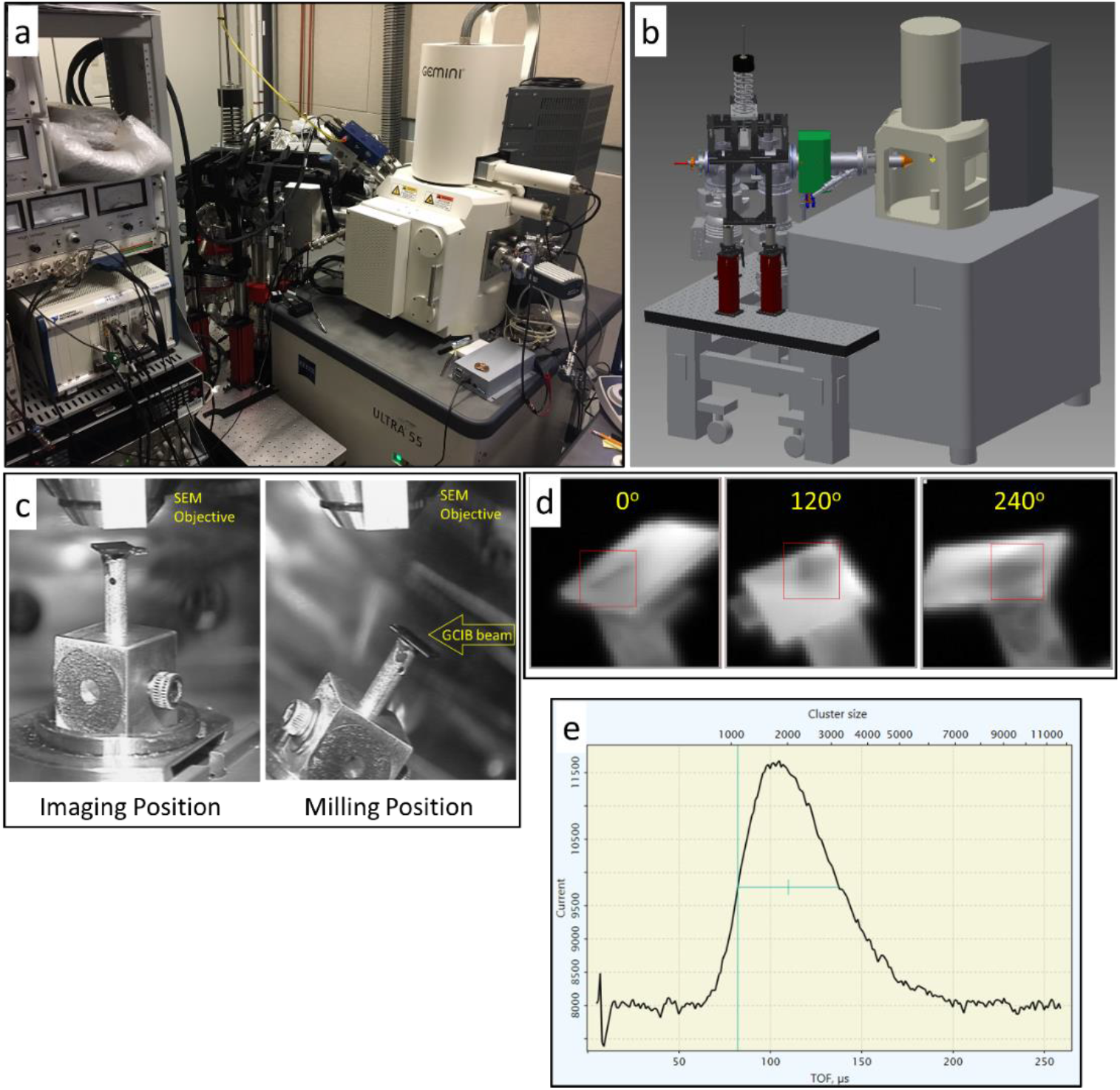
GCIB-SEM prototype constructed at the Janelia Research Campus. **(a)** Picture of the GCIB-SEM prototype consisting of an Ionoptika GCIB-10s gun mounted on a Zeiss Ultra SEM. Ion beam and SEM scanning and imaging are performed using electronics from National Instruments (rack visible in picture) and controlled via custom LabVIEW software. **(b)** CAD diagram showing how GCIB-10s gun was mounted. The considerable weight of the GCIB-10s gun, along with its two vibration isolated turbo pumps, was hung via a spring-loaded rod from a mount which is itself supported on an L-shaped vibration isolation table. **(c)** Two views (at imaging position and at milling position) of a sample within the SEM chamber (taken by an in-chamber CCD camera mounted at the back of the chamber). Sample is a series of thick tissue sections collected on a small (5 x 7 mm) silicon wafer. The wafer was mounted on the end of a short tube in order to make its silhouette visible in GCIB-scanned images. **(d)** GCIB-scanned overview images of sample acquired by measuring the sample current during GCIB scanning. Silhouette of wafer is clearly visible in these overview images which assists the targeting of the GCIB milling box (red square) used during the run. We rotate the sample to different milling directions (e.g. 0°, 120°, 240°) during each milling cycle as shown here. **(e)** Plot showing the range of cluster sizes hitting the sample (GCIB setting: Ar2000, 10kV). This is a time-of-flight plot made by pulsing the GCIB source while the beam is hitting a sample mounted at a known distance. The GCIB was tuned to give clusters containing an average of 2000 argon atoms each, but the plot shows that this beam actually contains a range of cluster sizes from mainly within the range of Ar1000 to Ar3000.

**SOM Figure 2.**
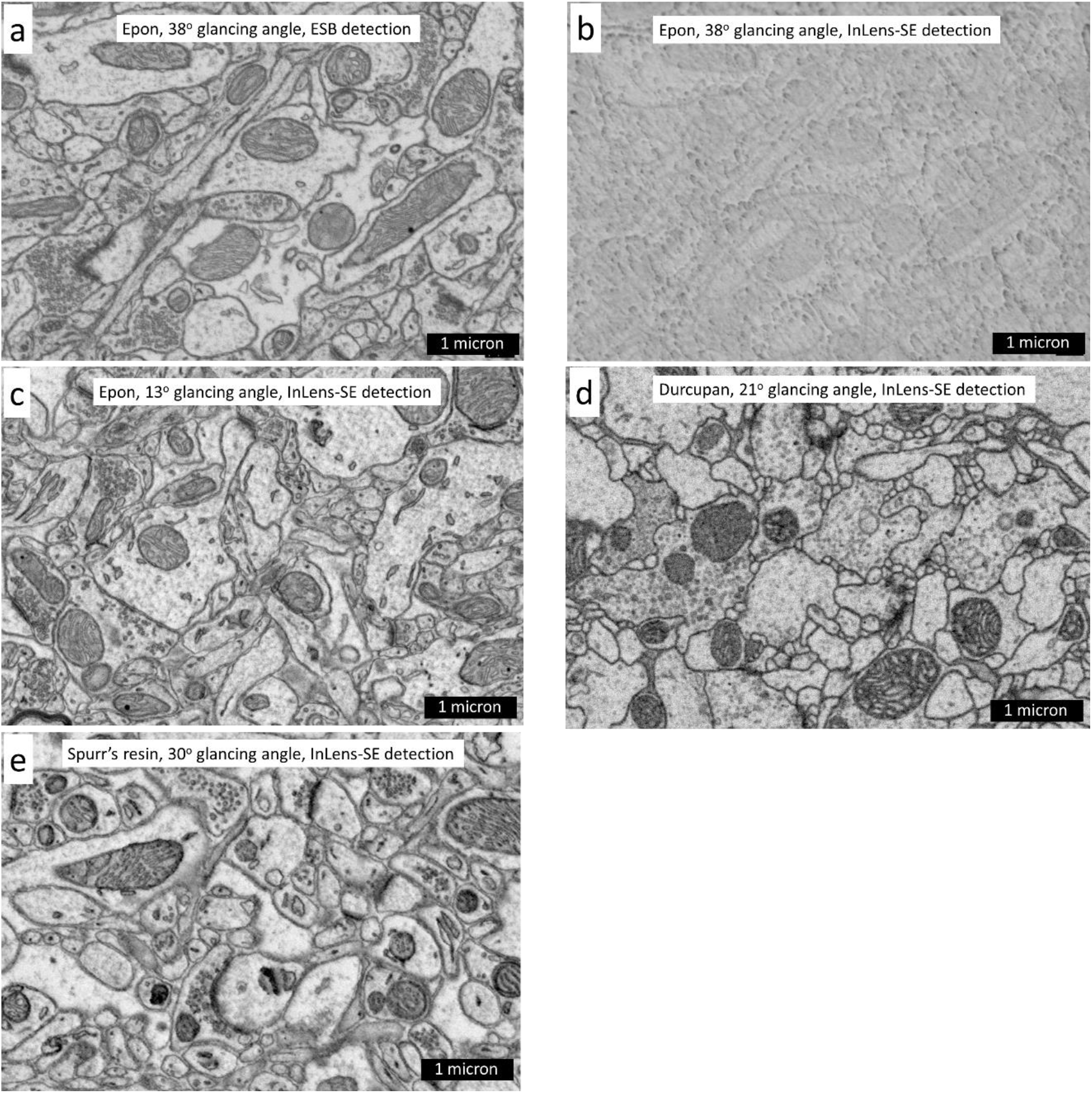
ESB and InLens-SE images of tissue surfaces GCIB-milled under different conditions and embedded in different resins. 1.2kV landing energy was used for all. 500v energy filter used in ESB. **(a)** ESB image of Epon-embedded section of mouse cortex milled using Ar2000, 10kV at a glancing angle of 38°. **(b)** InLens-SE image of same surface shown in **(a)**. This matched pair of images dramatically illustrates how much more sensitive InLens-SE detection is to surface topography. **(c)** InLens-SE image of epon-embedded section of mouse cortex milled using Ar2000, 10kV at a glancing angle of 13°. This shallow angle milling has succeeded in removing the InLens-SE-visible surface structure seen in **(b)** but at a much reduced milling rate. **(d)** InLens-SE image of Durcupan-embedded section of *Drosophila* brain milled using Ar2000, 10kV at a glancing angle of 21°. **(e)** InLens-SE image of Spurr’s-embedded section of mouse cortex milled using Ar2000, 10kV at a glancing angle of 30°.

**SOM Figure 3.**
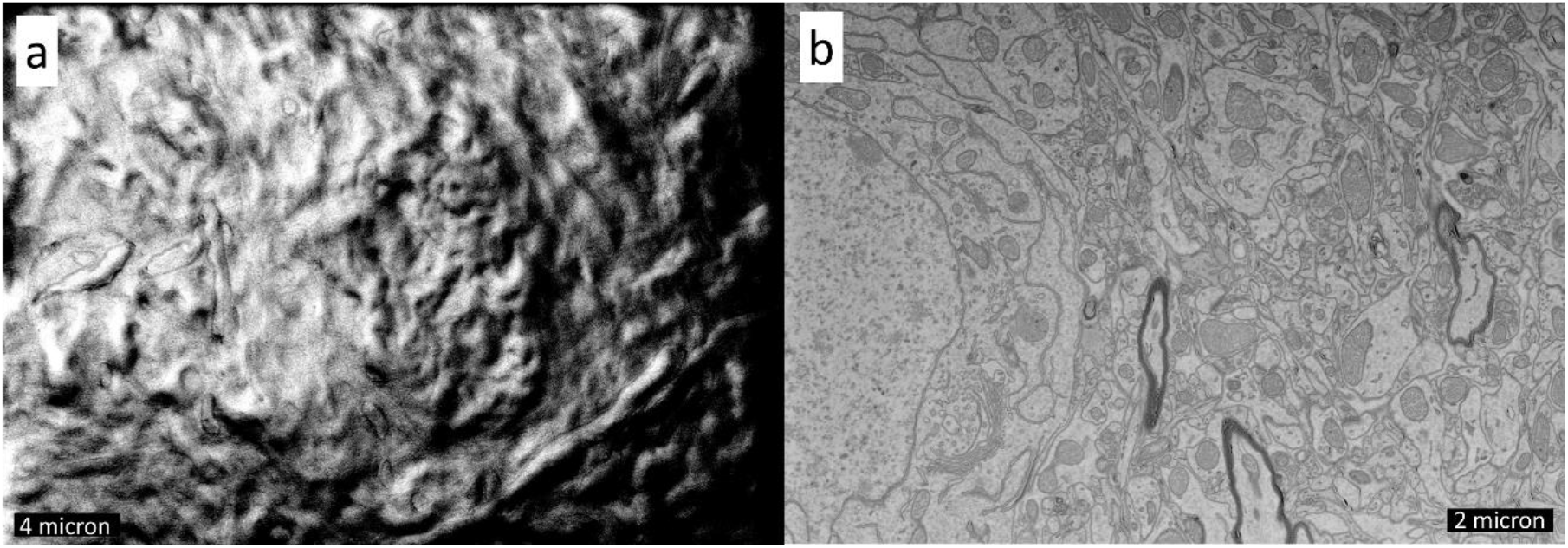
Using electron irradiation to increase a sample’s bulk conductivity. **(a)** InLens-SE image of a 400 nm thick section of plastic-embedded mouse cortex imaged at 1.2 kV landing energy. Sample charging results in large intensity and scan distortions making imaging impossible. **(b)** Same 400 nm thick section after being electron irradiated with an 8 kV beam to increase its bulk conductivity. InLens-SE imaging at 1.2 kV landing energy now gives clear images of neuronal features.

**SOM Figure 4.**
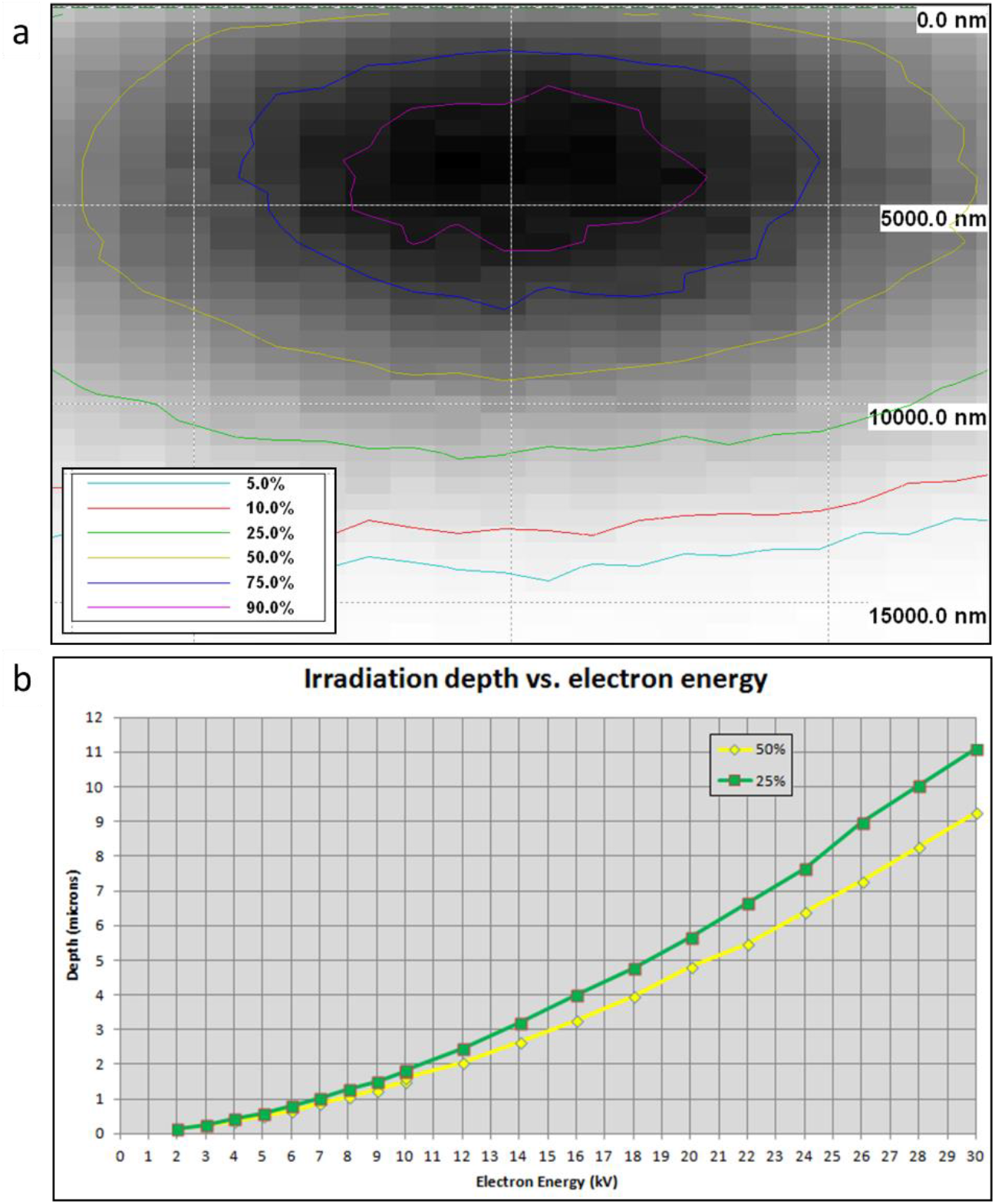
Estimation of optimal electron energy for irradiation. **(a)** Energy density plot, from the Casino ‘electron trajectories in solids’ simulator (Drouin et al. 2007), of a wide beam of 30 kV electrons irradiating an Epon block with 2% Osmium atomic fraction. Yellow contour line shows where the energy density has dropped to 50% of maximum, green contour line shows where the energy density has dropped to 25% of maximum. **(b)** Summary plot of Casino simulations covering 2kV to 30kV beam energies. Yellow and green points show depths where energy density drops to 50% and 25% of maximum respectively. We use this plot to determine what beam energy should be used for irradiating a sample of a given thickness.

**SOM Figure 5.**
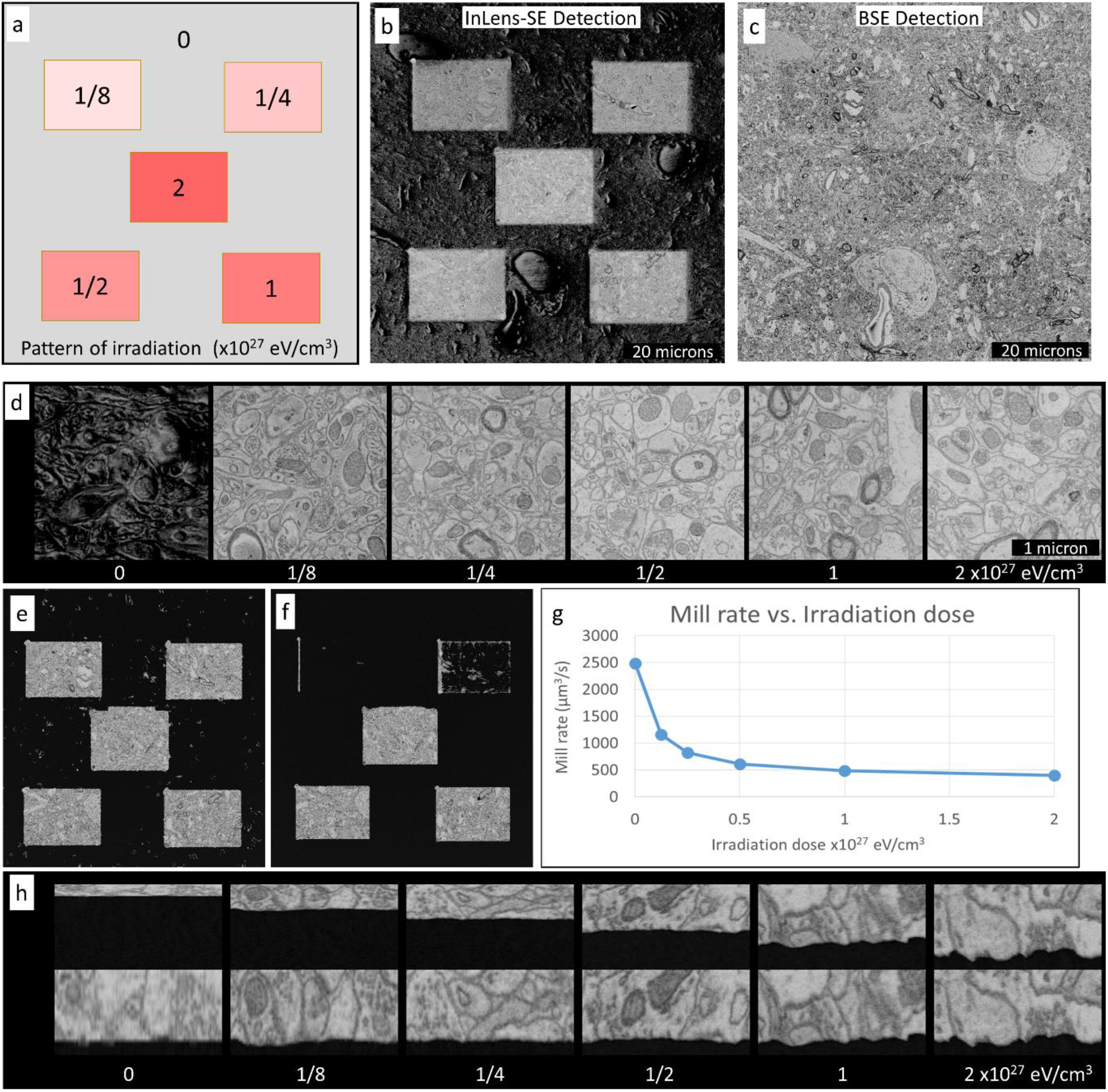
Imaging and milling rate tests conducted on a 500 nm thick section of Spurr’s embedded mouse cortex irradiated under a range of conditions. **(a)** Pattern of electron irradiation used for test. Five rectangles, each 20×15 μm, were electron irradiated with a defocused 6 kV electron beam for varying durations to achieve the following total doses: 0.13×10^27^, 0.25×10^27^, 0.50×10^27^, 1.0×10^27^, 2.0×10^27^ eV/cm^3^. Surrounding tissue was not irradiated beyond that induced by imaging. **(b)** InLens-SE detector image taken after five mill cycles. Charging artifacts are evident in non-irradiated and lightly-irradiated regions. **(c)** ESB detector image taken after five mill cycles. Only mild charging artifacts (e.g. the blood vessel at bottom center) are seen in this ESB image. SEM imaging parameters: 1.2 kV, 2 nA electron beam, 10 nm pixels, 1.25 MHz acquisition rate. **(d)** Zoomed-in InLens-SE images of regions in **(a)** showing extent of charging. Non-irradiated regions of this 500 nm section show massive charging artifacts in InLens-SE. The 0.13×10^27^, 0.25×10^27^ eV/cm^3^ irradiated regions show milder charging artifacts which appear as contrast differences within mid-sized regions of cytoplasm. Regions which received greater irradiation show no charging artifacts in InLens-SE. **(e)** Image of tissue after 17 milling cycles. All non-irradiated regions have been milled away. **(f)** Image of tissue after 44 milling cycles. The two least-irradiated regions have also been milled away at this point in the run. Milling conditions: 32nA of Ar2000 at 10 kV spread over a 12.5 mm^2^ area, 30° glancing angle, 360° rotation (3 steps), 180 s per mill cycle. **(g)** Plot of milling rate vs. irradiation dose. There is a six fold drop in milling rate between the non-irradiated region and the most heavily irradiated region. **(h)** Top row shows z-reslice through aligned, but otherwise unmodified, ESB image stacks at each of the different irradiation doses. All of these extend through the entire 500 nm depth of the section but have different voxel sizes along the z-axis due to their different milling rates. For example, the stack through the non-irradiated region milled through to the gold substrate after only 14 mill cycles giving a voxel size of 10×10×36 nm (i.e. 500 nm/14 = 36 nm). In contrast, the most heavily irradiated regions milled through to the gold substrate after 86 milling cycles giving a voxel size of 10×10×5.8 nm (i.e. 500 nm/86 = 5.8 nm). Bottom row shows same z-reslice views after each was scaled in Z to have matching voxel sizes. This scaling makes clear that higher irradiation dose also results in increased local variability of milling rate as evidenced by the greater unevenness at the tissue-to-substrate boundary at the bottom of the most heavily irradiated region’s stack. Based on these results, we recommend irradiating to a dose of 0.5×10^27^ eV/cm^3^.

**SOM Figure 6.**
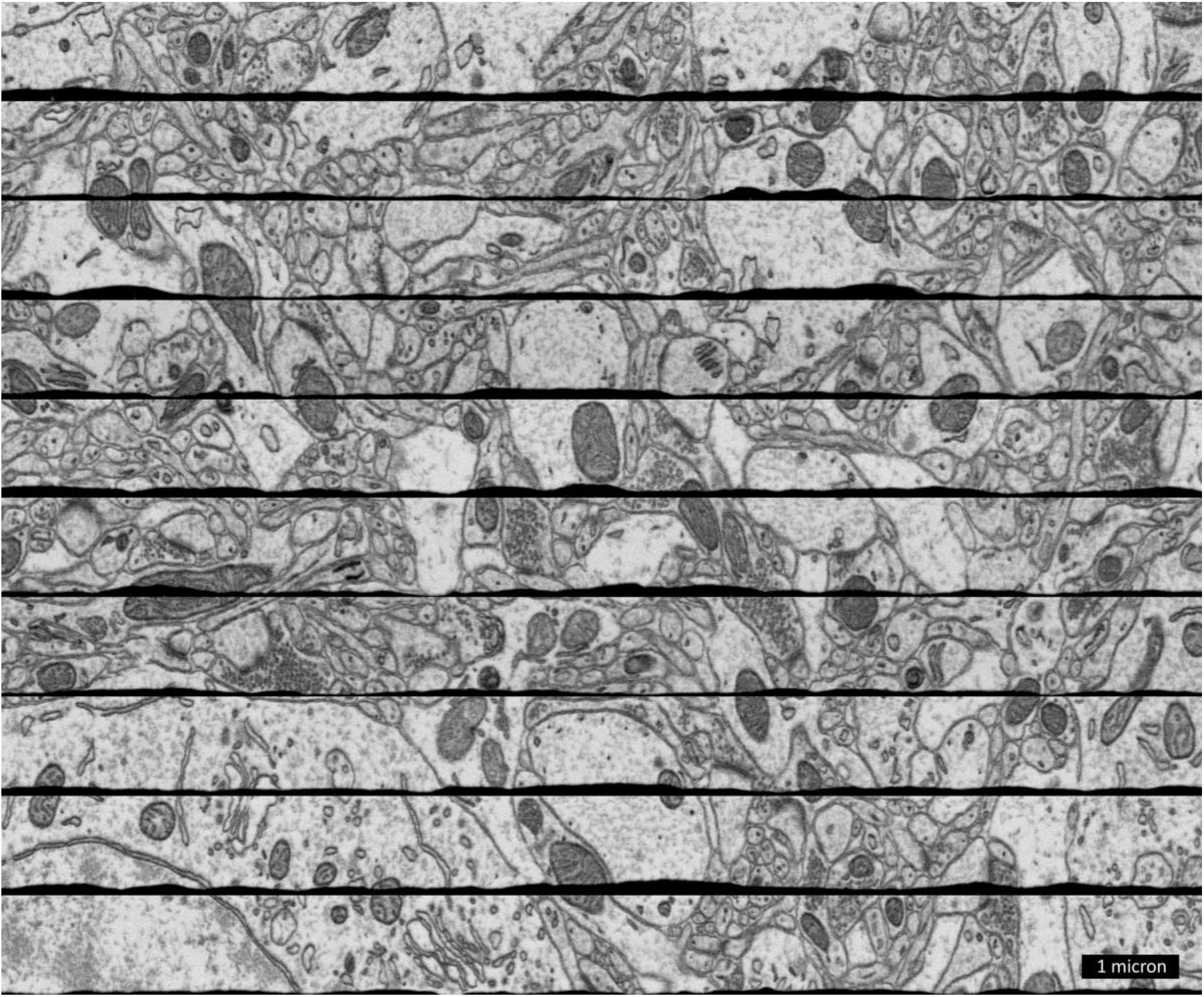
Z-reslice of GCIB-SEM stack covering ten 1 μm thick sections of Spurr’s-embedded mouse cortex (same dataset as shown in **Fig. 2d-f**). The GCIB-SEM stacks of the ten separate sections have been individually aligned and then aligned to each other to make the above stack, but the individual stacks have not yet been flattened giving rise to the black unevenness at the bottom of each section. (ESB detection)

**SOM Figure 7.**
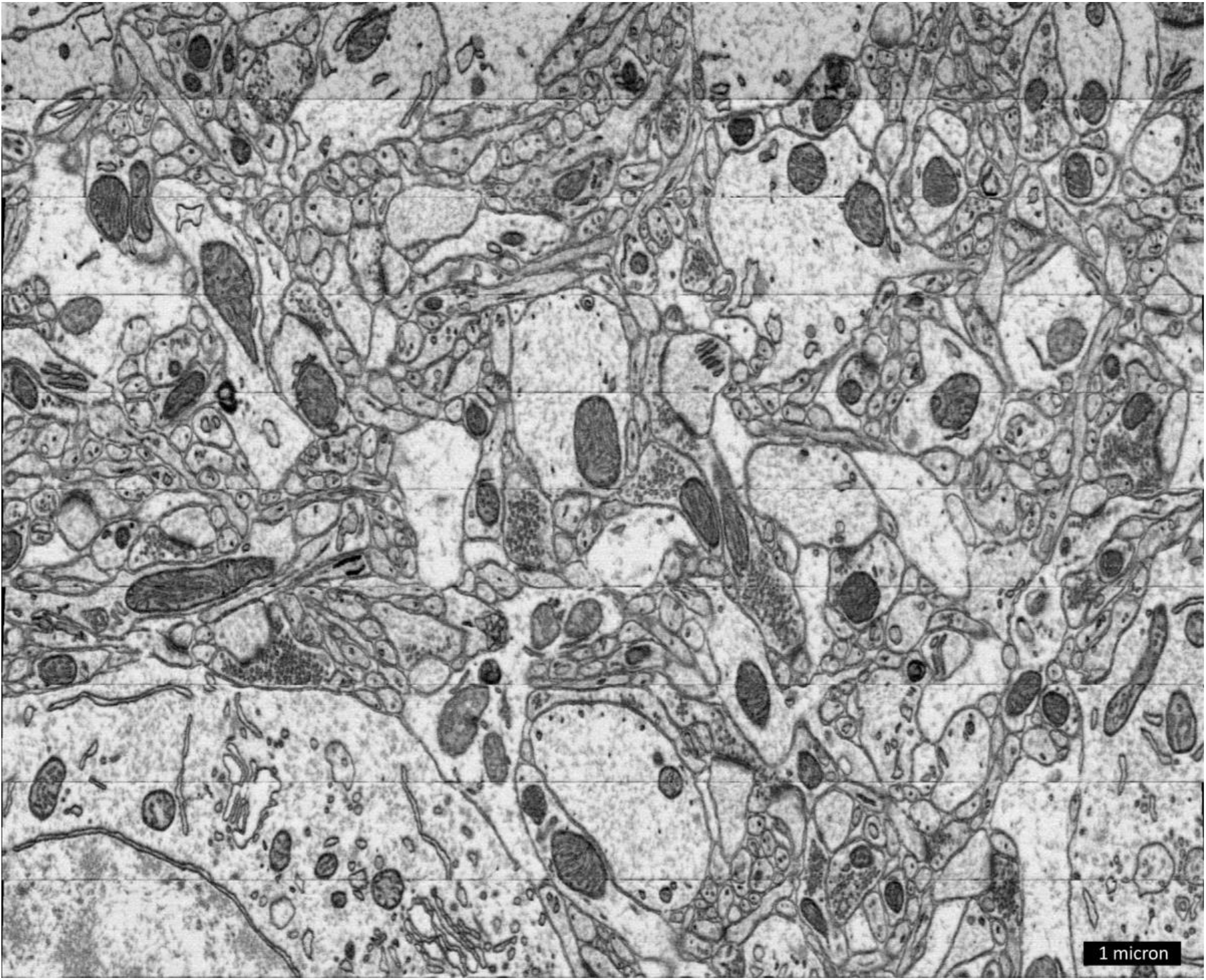
Same GCIB-SEM stack as shown in the previous figure but after computationally flattening each of the ten 1 μm thick sections and then stitching them together. This flattened and stitched volume is the one that was segmented and traced as shown in **Fig. 2e.** (ESB detection)

**SOM Figure 8.**
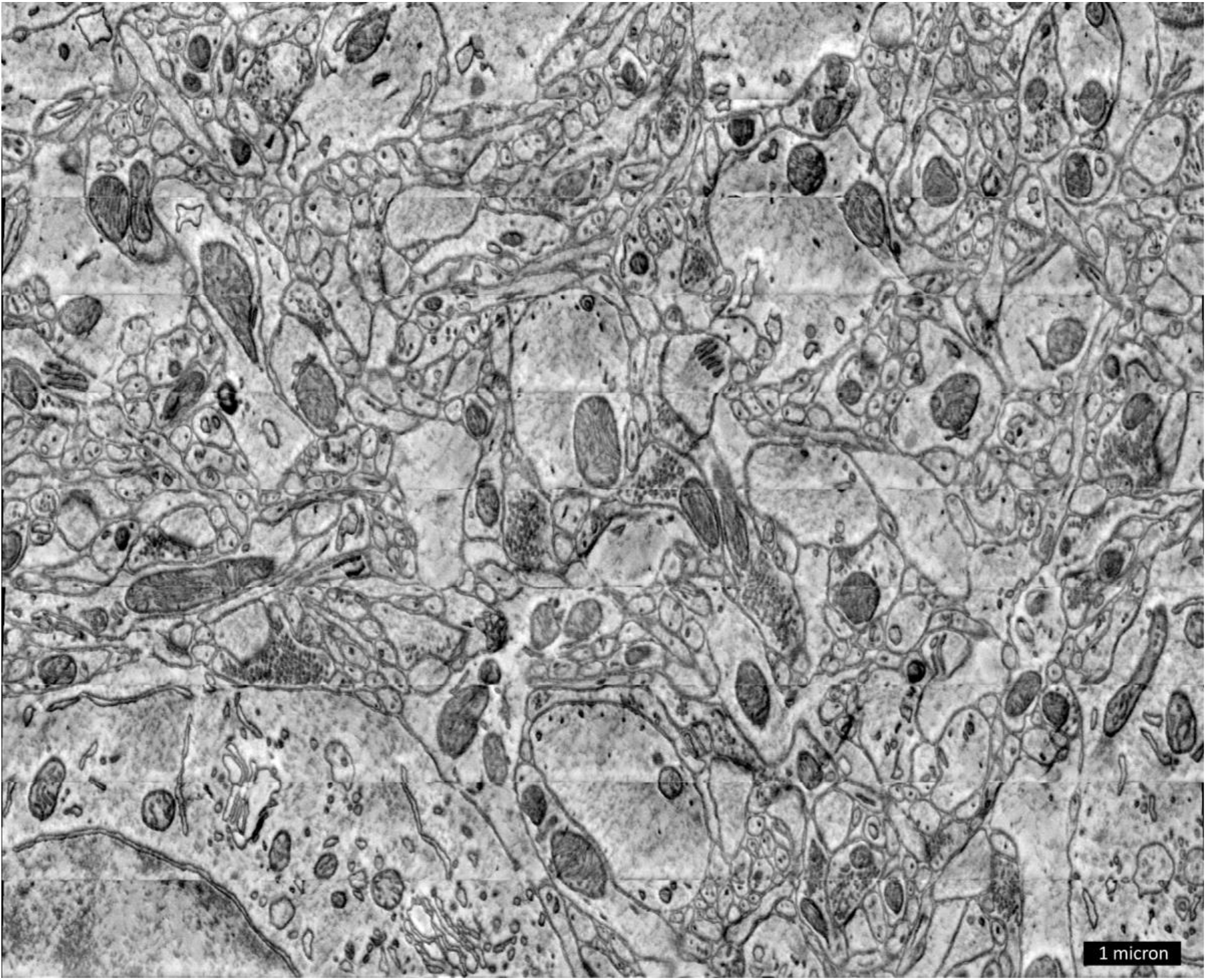
Same GCIB-SEM volume as shown in the previous figure but based on the InLens-SE detector signal. Slight charging artifacts are visible as intensity variations within mid-sized cytoplasm regions. This sample was irradiated with an approximately 2.4×10^26^ eV/cm^3^ dose. Such charging can be completely eliminated with larger irradiation dose (for example, see **SOM Figure 14** below).

**SOM Figure 9.**
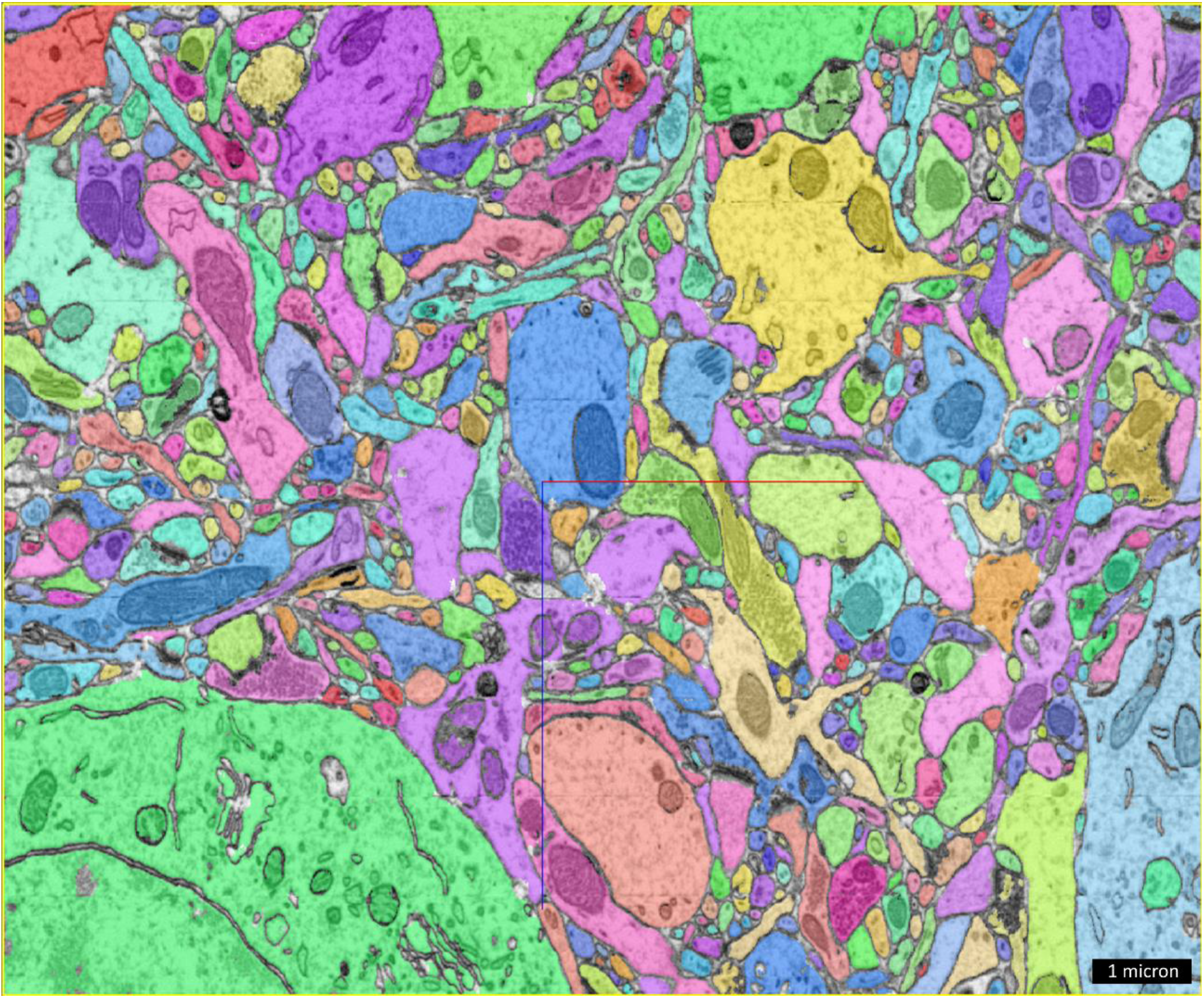
Z-reslice of the same GCIB-SEM volume as shown in the previous figures (and in **Fig. 2d-f**) segmented using a flood-filling network (Januszewski et al. 2018).

**SOM Figure 10.**
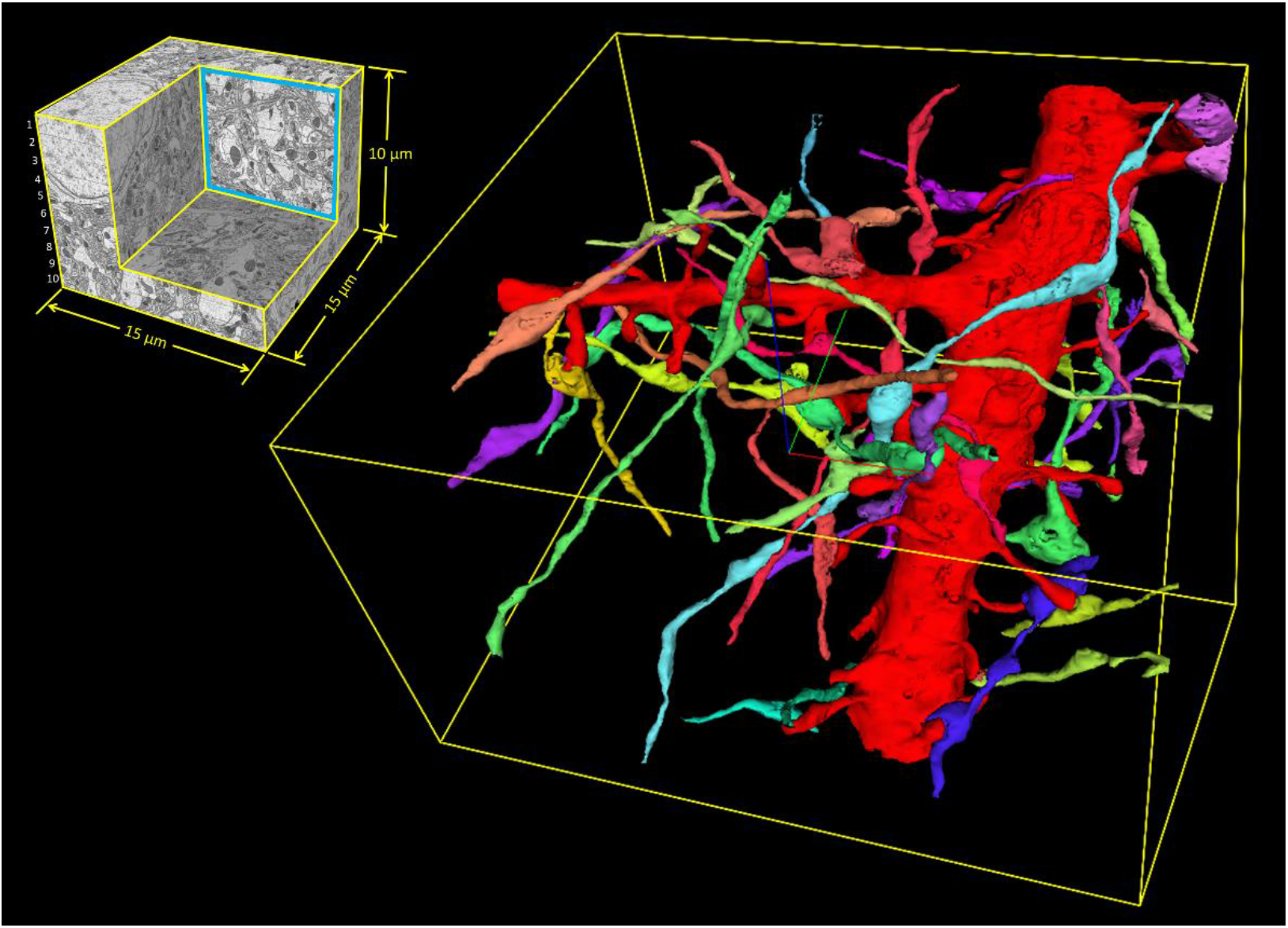
Example tracing, within the same GCIB-SEM volume as shown in the previous figures, of all axons making synapses on a large spiny dendrite (red) in the volume.

**SOM Figure 11.**
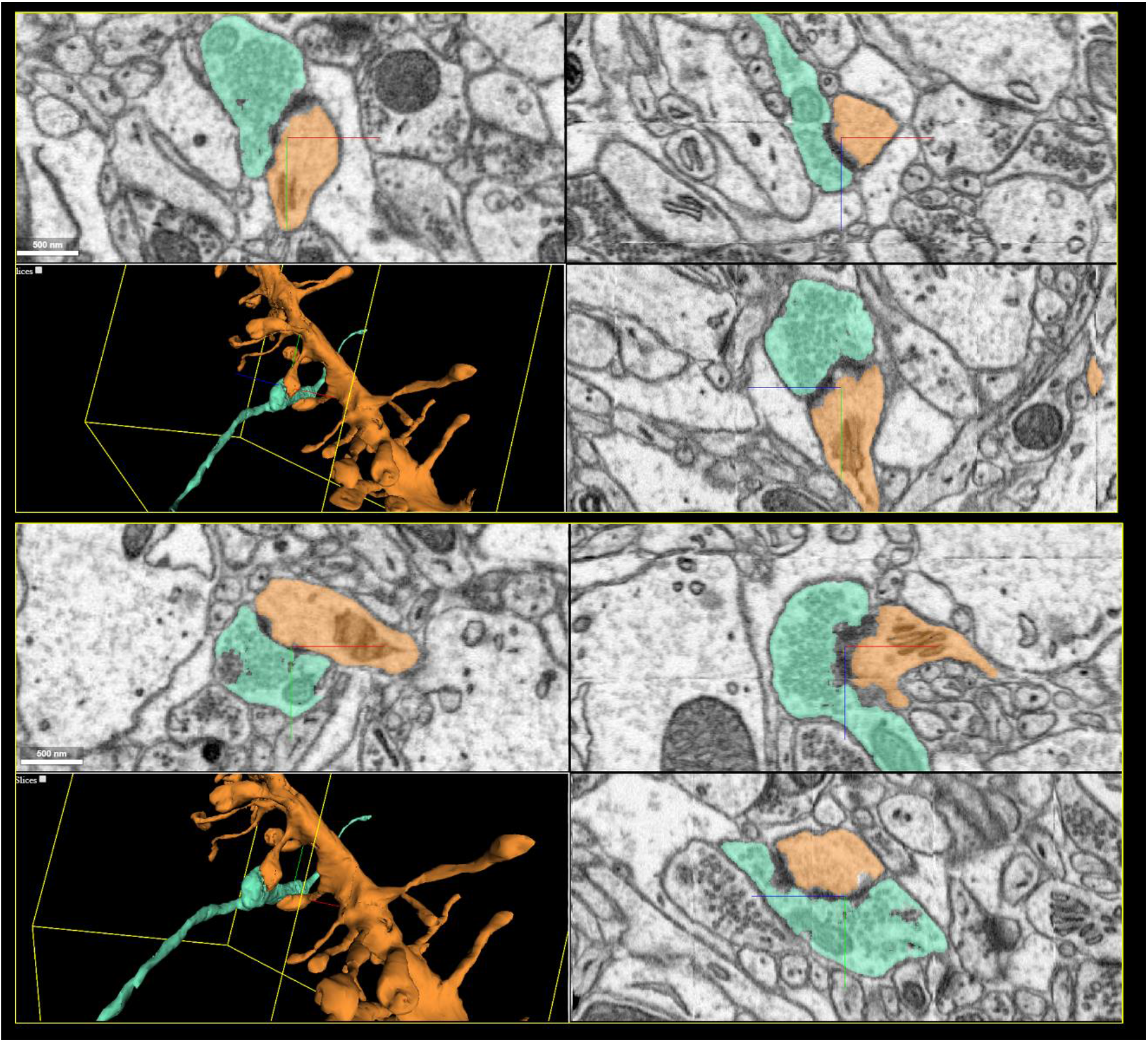
Example tracing, within the same GCIB-SEM volume as shown in the previous figures, of an axon (cyan) making two synapses on the same dendrite (orange).

**SOM Figure 12.**
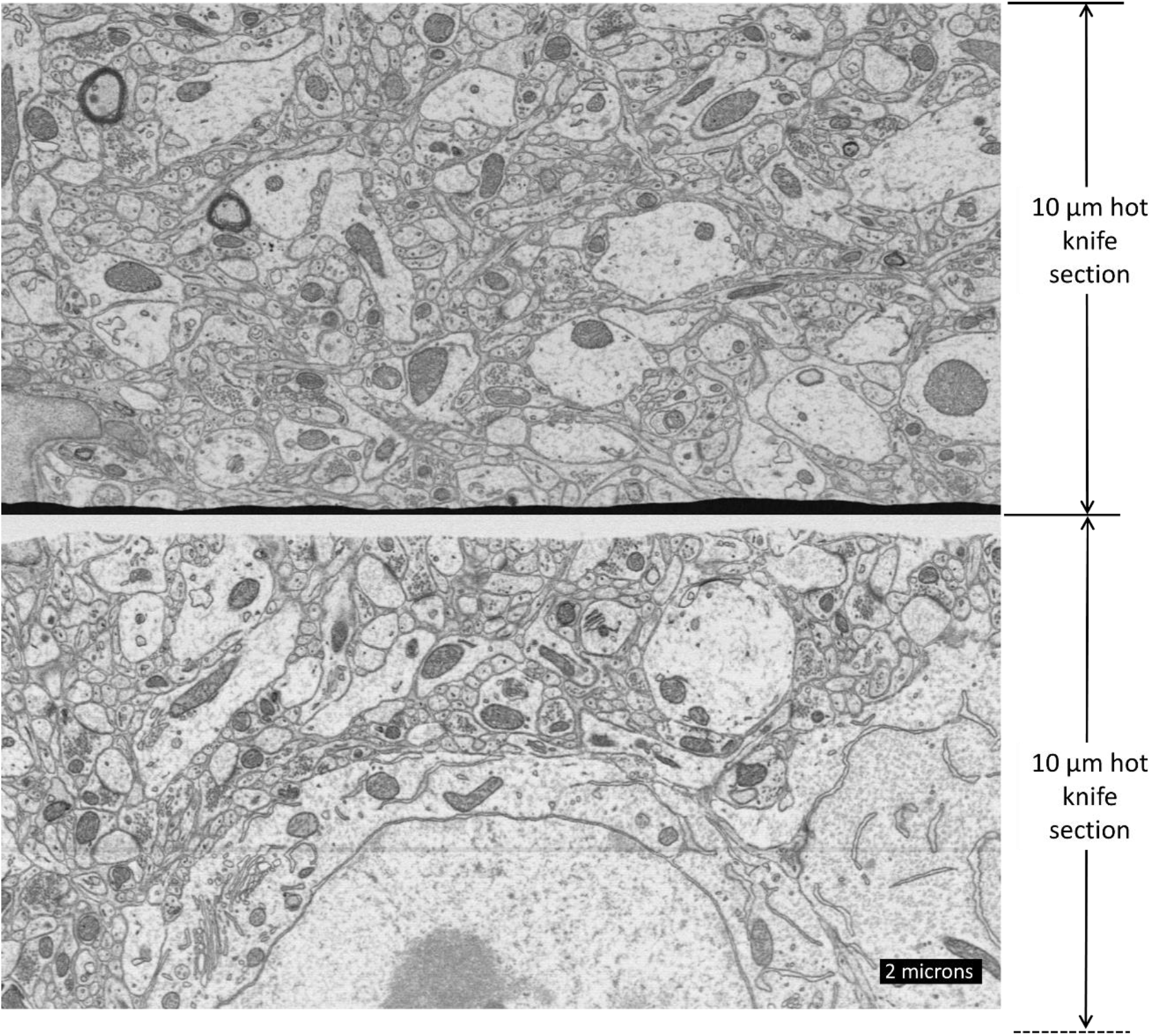
Z-reslice of GCIB-SEM stack covering two 10 μm thick hot knife sections of Spurr’s-embedded mouse cortex tissue (same dataset as shown in **Fig. 3**). The GCIB-SEM stacks of the two separate sections have been individually aligned and then aligned to each other to make the above stack, but the individual stacks have not yet been flattened. The black unevenness at the bottom of the top hot knife section is the gold substrate the section is embedded against. The unevenness at the top of the bottom section is resin covering the re-embedded section. (ESB detection)

**SOM Figure 13.**
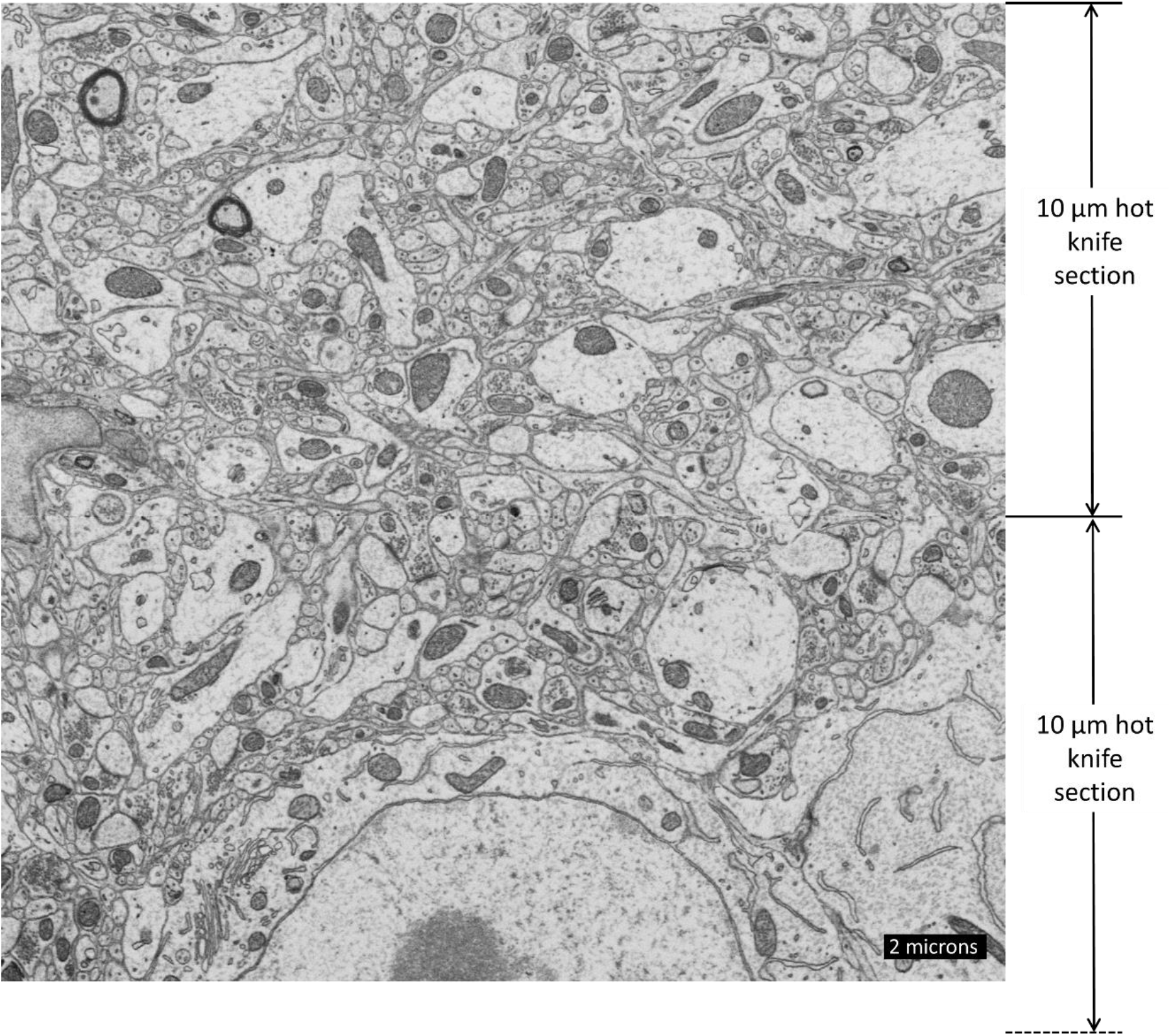
Same GCIB-SEM stack as shown in the previous figure after computationally flattening the bottom of the top hot knife section and after flattening the top of the bottom hot knife section. The location of the stitch line between the two is designated at right as it is barely visible in this z-reslice image. (ESB detection)

**SOM Figure 14.**
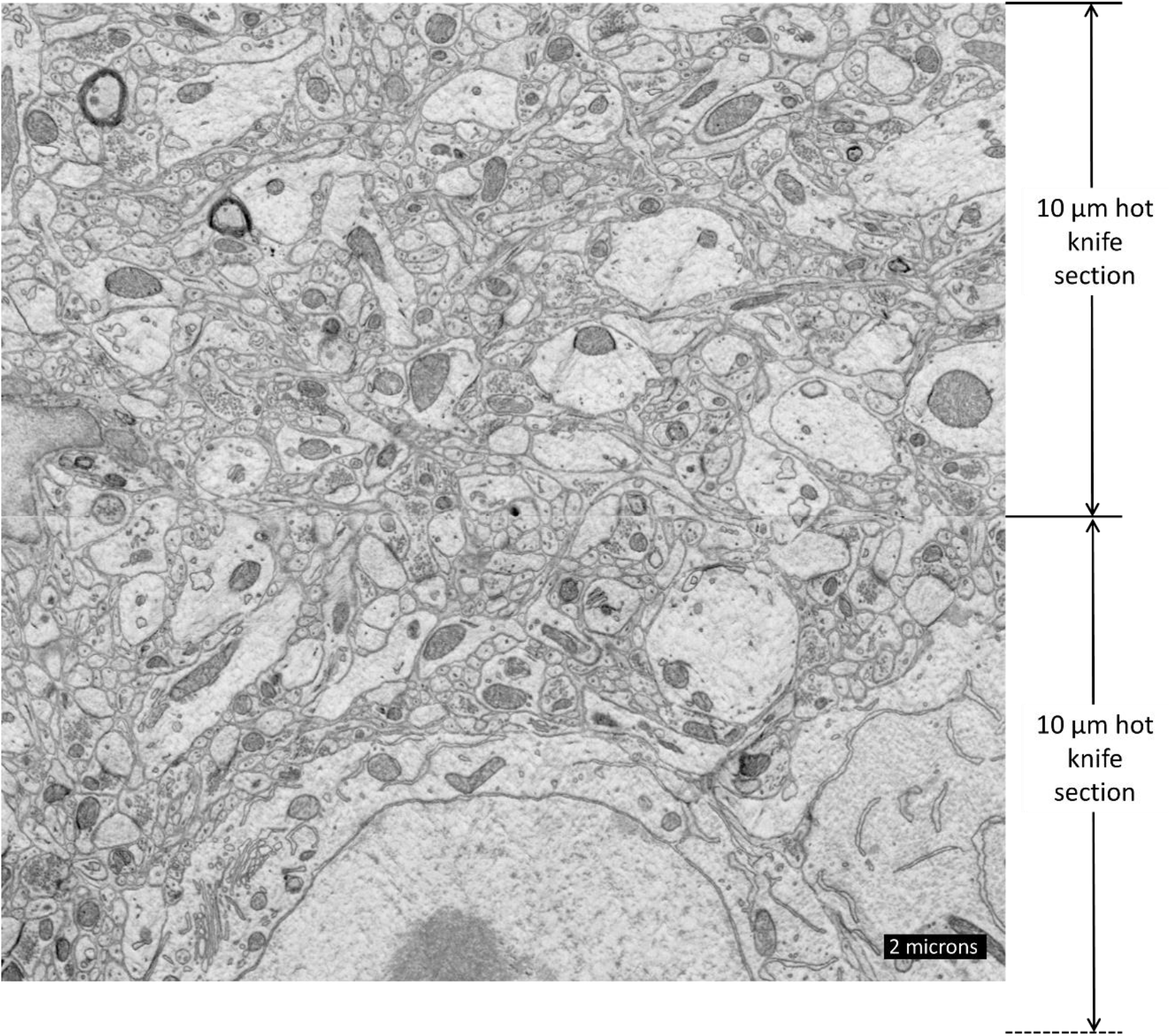
Same GCIB-SEM volume as shown in the previous figure but based on the InLens-SE detector signal. No charging artifacts are visible in this GCIB-SEM run due to the high electron irradiation dose used (0.9×10^27^ eV/cm^3^). Some streak milling artifacts are visible in this InLens-SE image that are not in the previous ESB image. The streak milling artifacts seem more prevalent at deeper milling depths. (InLens-SE detection)

**SOM Figure 15.**
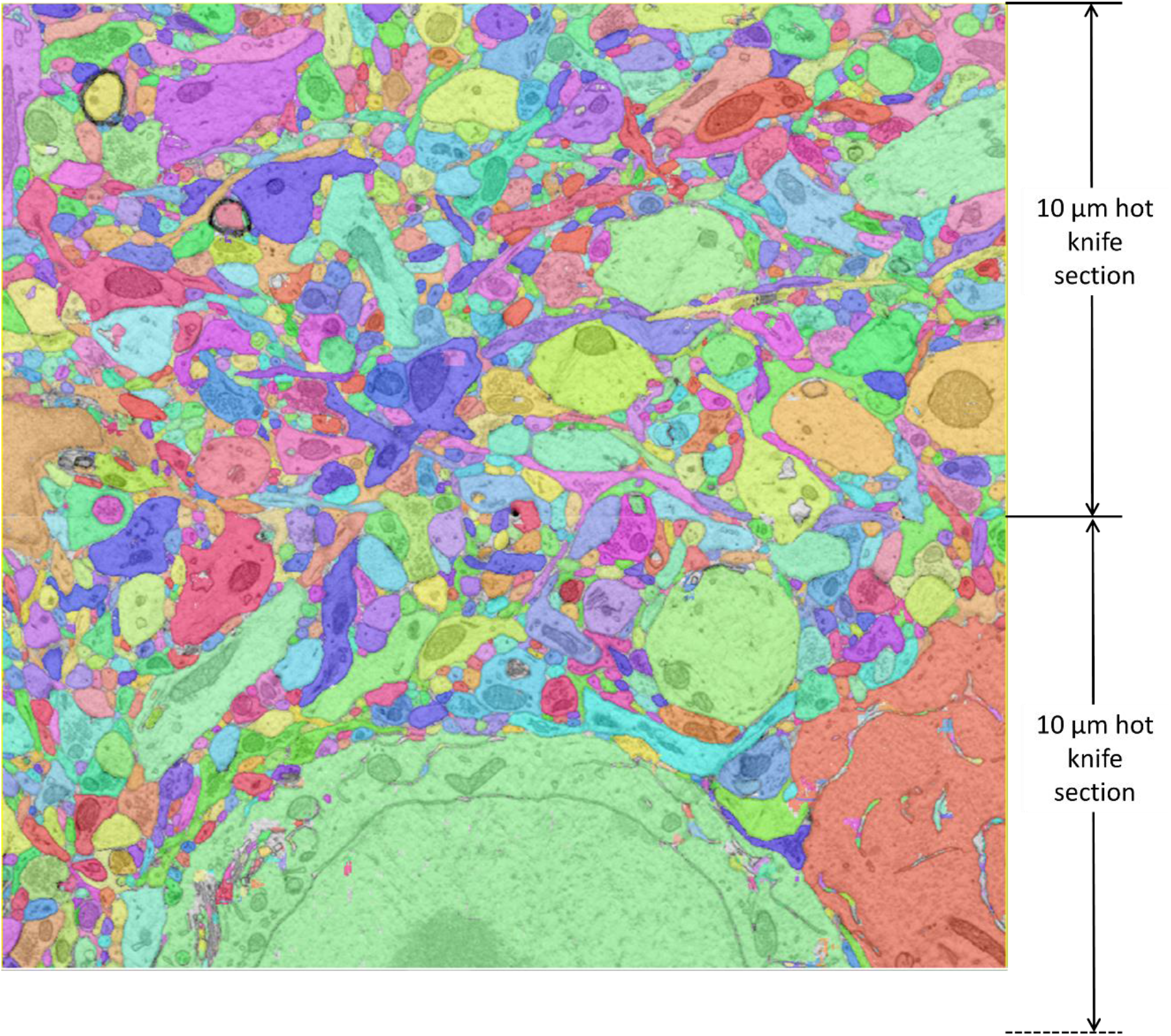
Same z-reslice as shown in previous image but with overlaid colors showing result of flood-filling network segmentation based on InLens-SE volume.

**SOM Figure 16.**
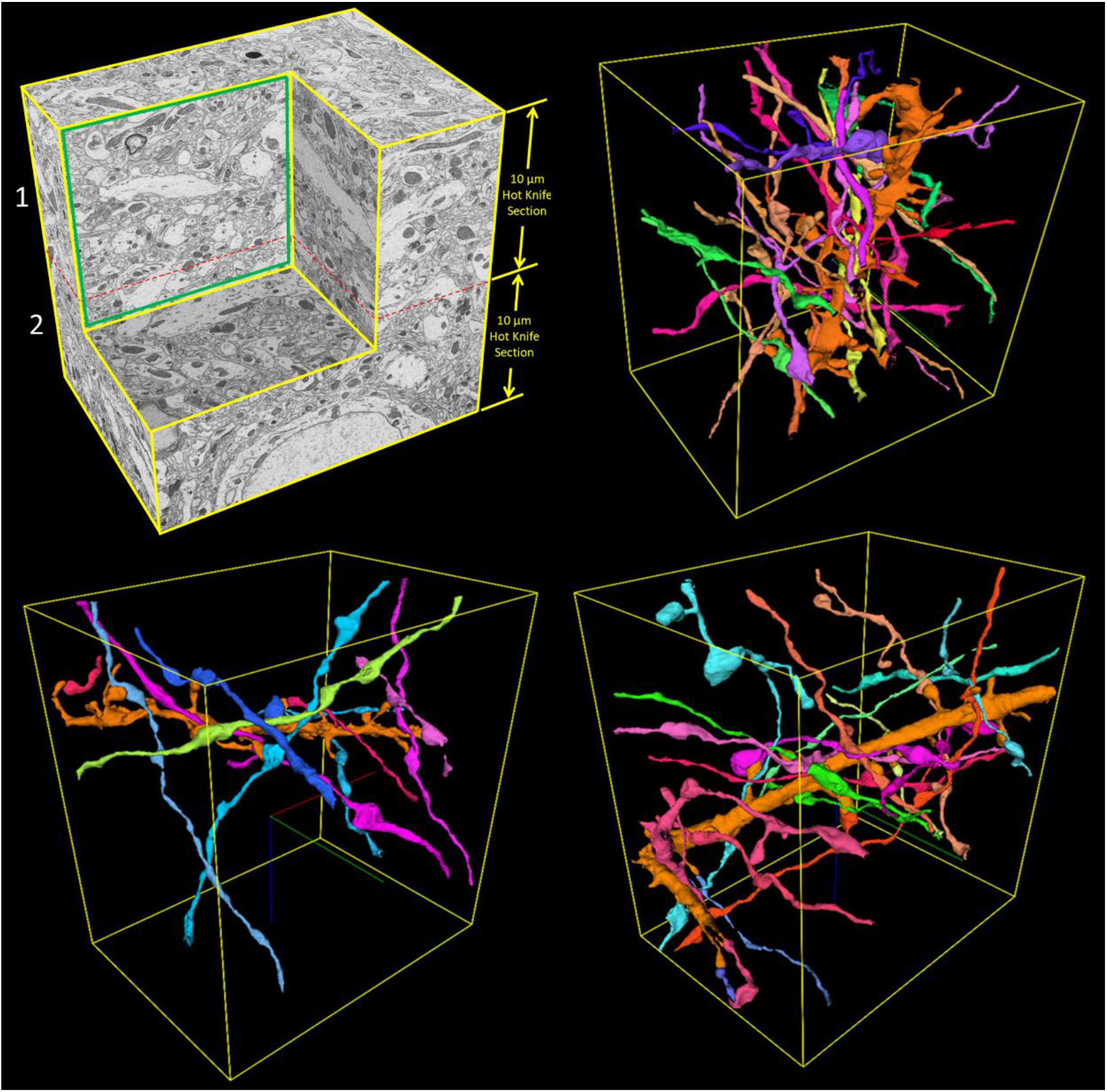
Example tracings, within the same flood-filling network-segmented GCIB-SEM volume as shown in the previous figure, of axons making synapses on three different dendrites (orange) spanning the volume.

**SOM Figure 17.**
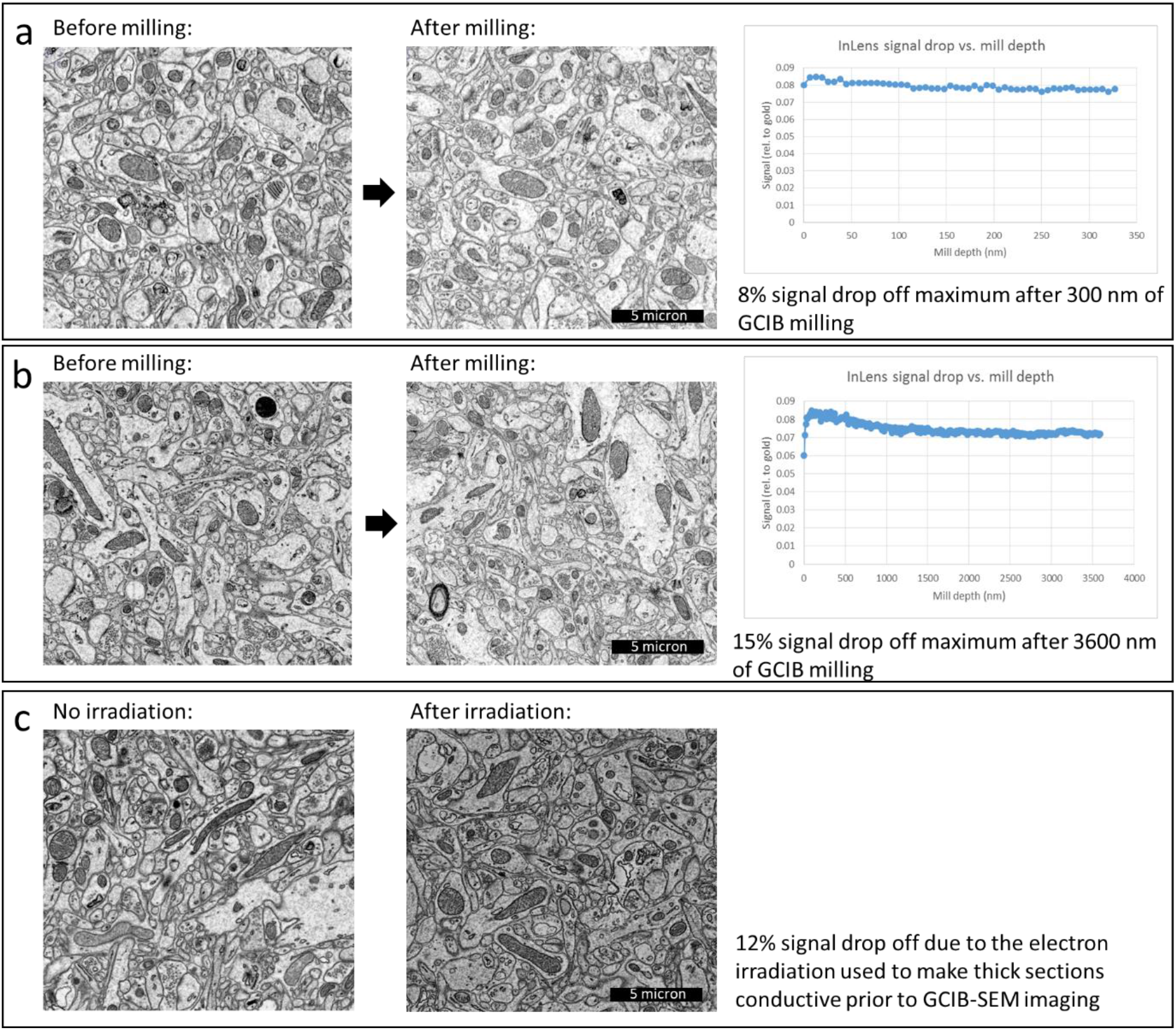
Estimates of InLens-SE signal loss (i.e. tissue contrast reduction) due to GCIB milling (a-b) and due to the electron irradiation used to make thick sections conductive prior to GCIB-SEM imaging **(c)**. **(a)** is from run shown in **Fig. 2a-c**. **(b)** is from run shown in **Fig. 3**.

**SOM Figure 18.**
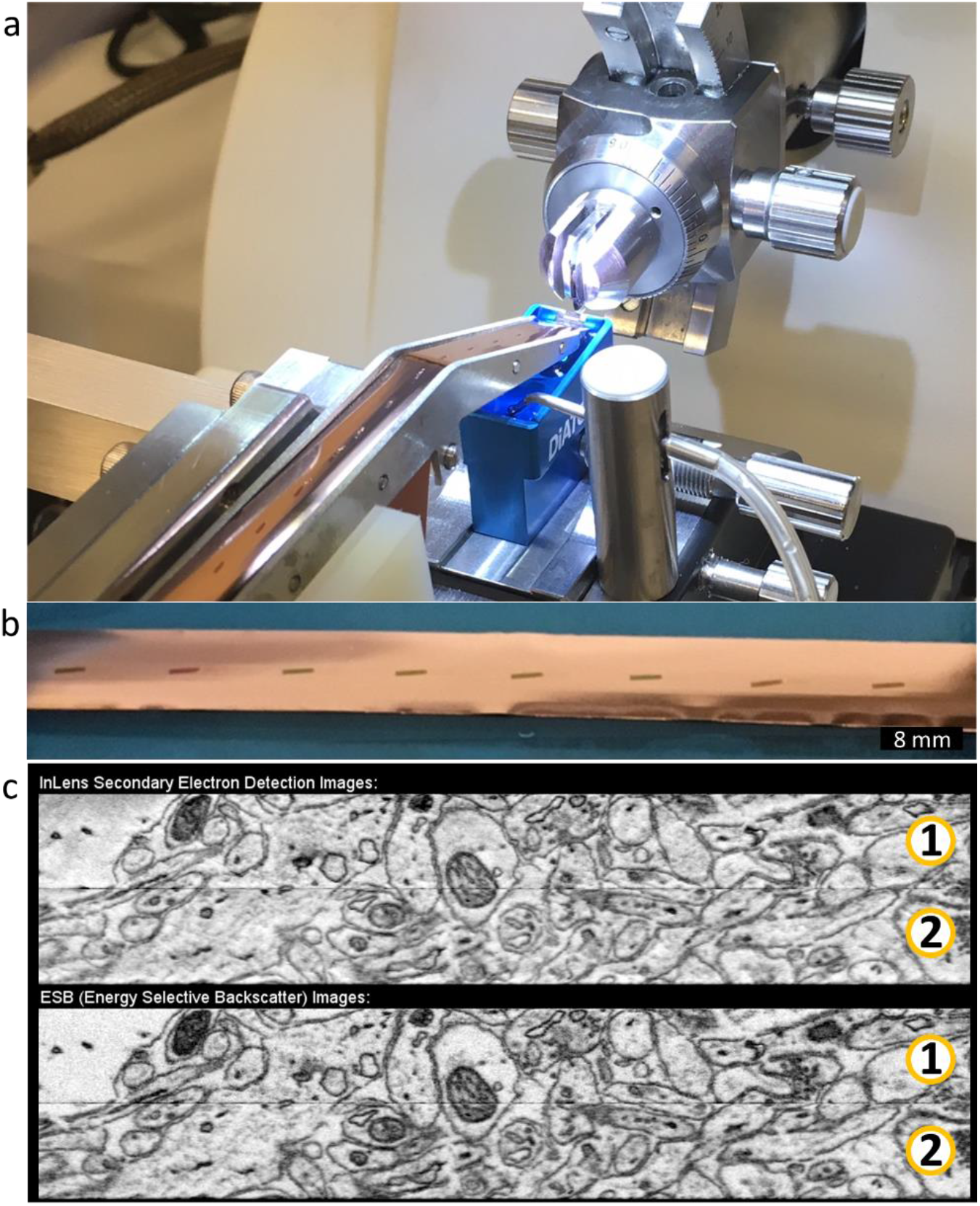
GCIB-SEM imaging of ATUM-collected sections. **(a)** Image of ATUM tape collection mechanism in the process of collecting a sequence of 1 μm sections of Spurr’s-embedded mouse cortex tissue on copper-coated tape (tape provided by Yoshiyuki Kubota). **(b)** Close-up of 1 μm sections on tape. **(c)** Z-reslice view (InLens-SE and ESB) through the GCIB-SEM volume spanning two sequential 1 μm sections collected during the above ATUM run.

**SOM Figure 19.**
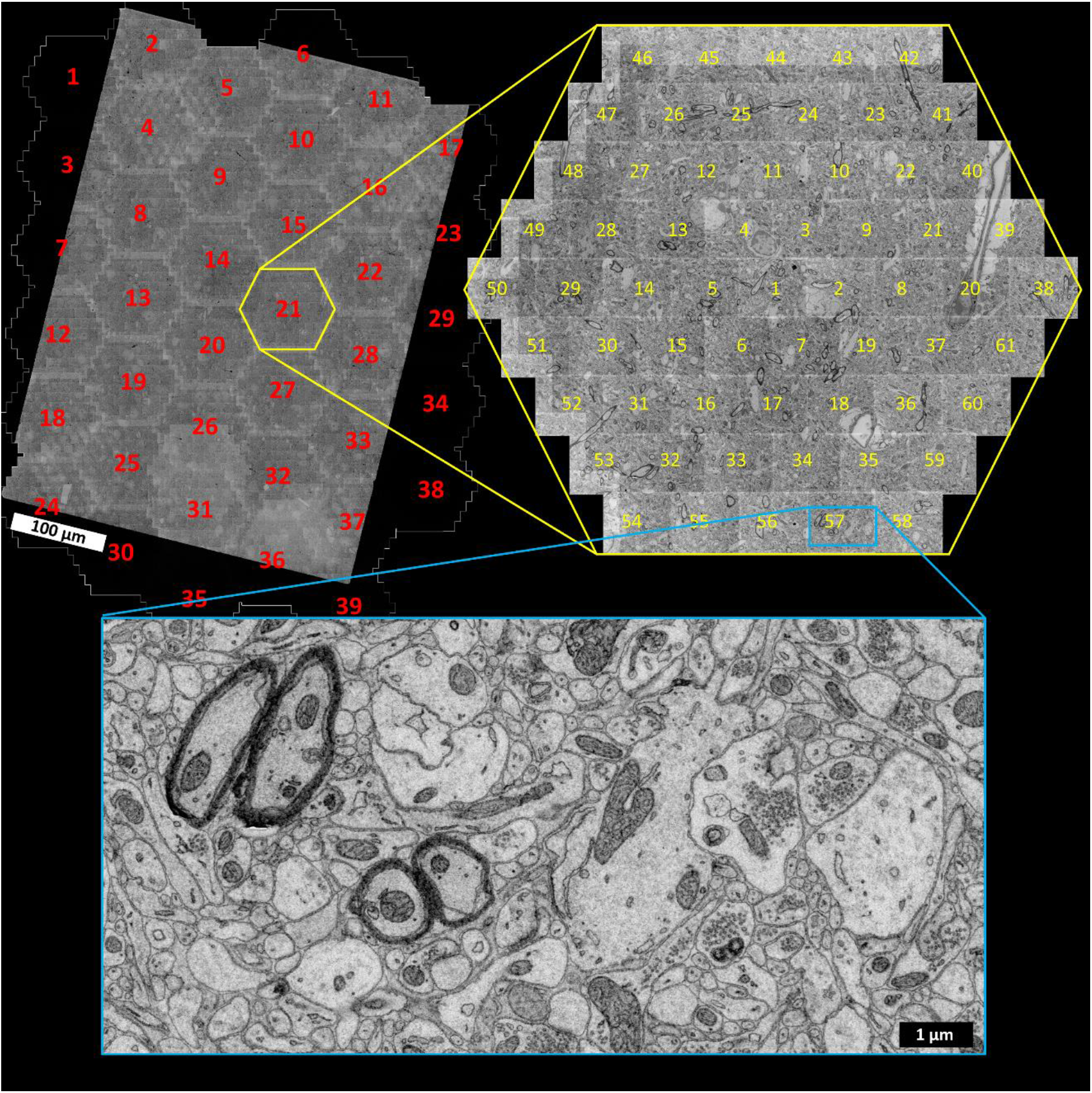
Test of Zeiss MultiSEM (61 beam version) imaging of a previously GCIB-SEM milled sample. A 500 nm thick section of Spurr’s-embedded mouse cortex tissue was electron irradiated to a dose of 0.9×10^27^ eV/cm^3^. It was then GCIB-SEM milled and imaged on our prototype GCIB-SEM at Janelia to a depth of ^~^250 nm. The sample was then shipped to Zeiss for imaging on the MultiSEM, the tiled images of which are shown in this figure. (Images acquired by Anna Eberle)

**SOM Figure 20.**
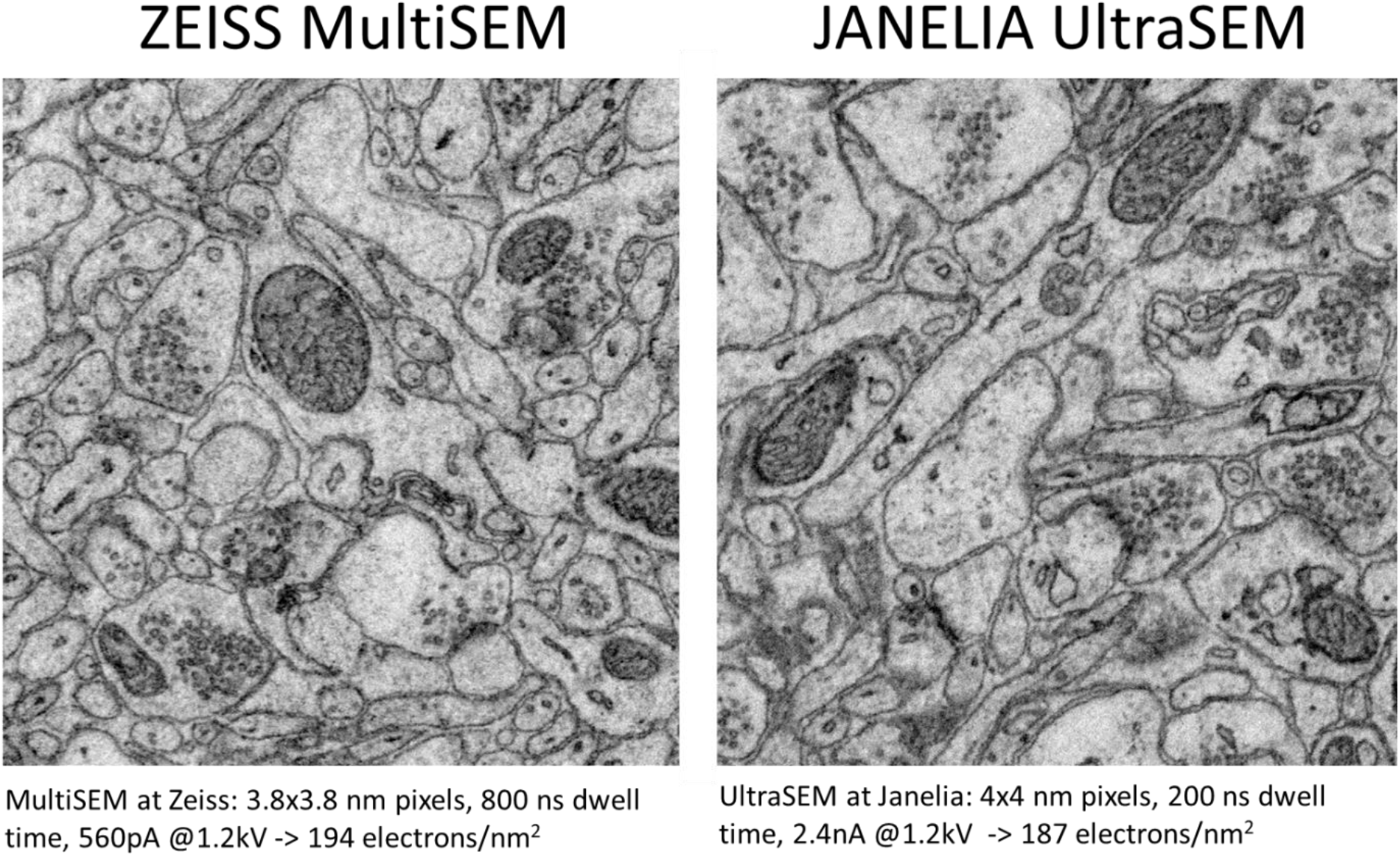
Comparison of Zeiss MultiSEM (61 beam version) imaging of a previously GCIB-SEM milled sample to an InLens-SE detected image taken in our prototype of the same GCIB-milled surface using similar dose.

**SOM Figure 21.**
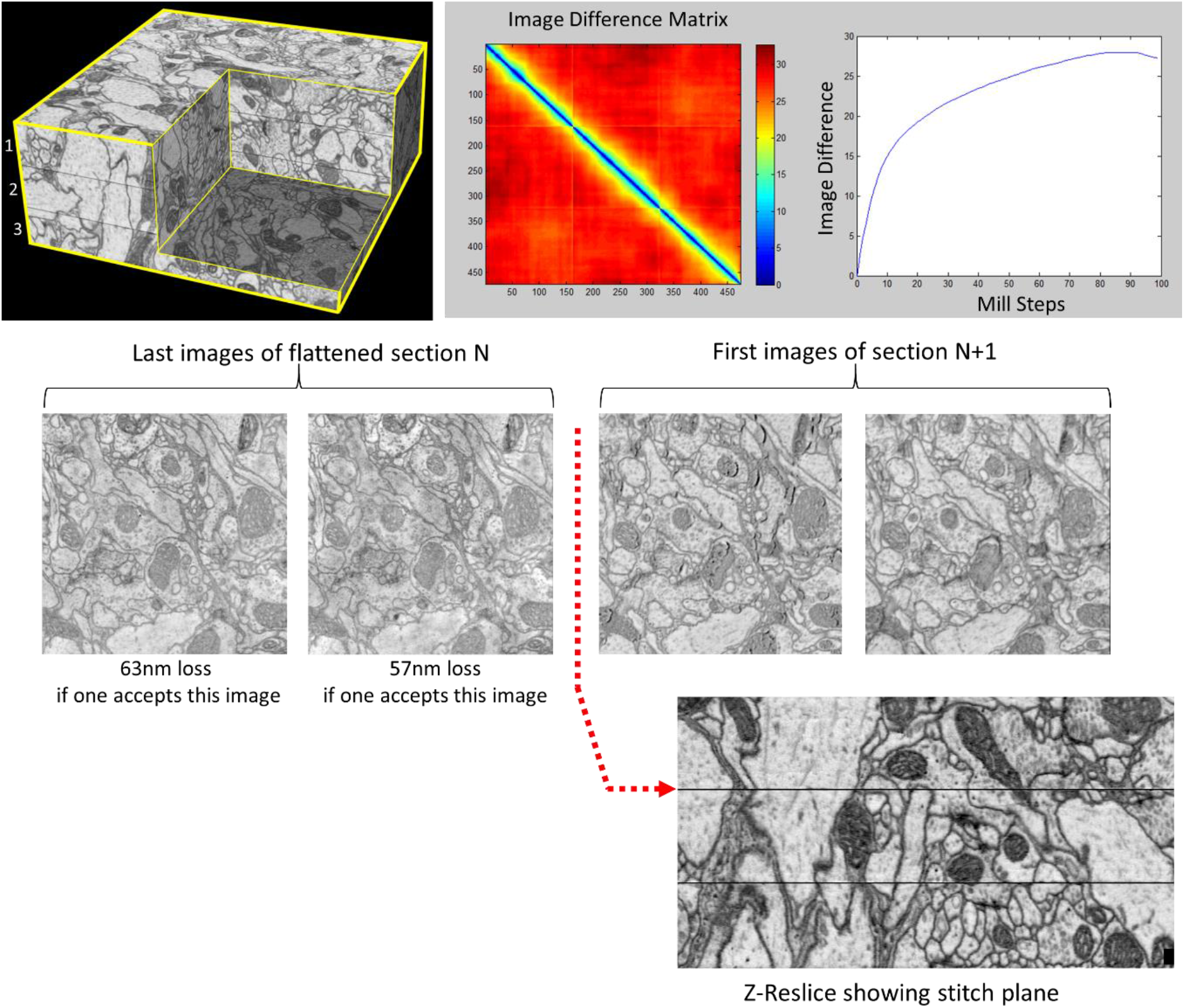
Estimate of sectioning loss for 1 μm thick sections of Durcupan-embedded fly brain tissue (same dataset shown in **Fig. 1b-e**). Note that the diamond knife cut surfaces show considerable knife damage, and the estimated loss between sequential sections is relatively large compared to the Spurr’s embedded samples below. This and other observations led us to conclude that Durcupan embedding is insufficiently resilient to be sectioned this thick. We therefore switched to Spurr’s embedding for later runs.

**SOM Figure 22.**
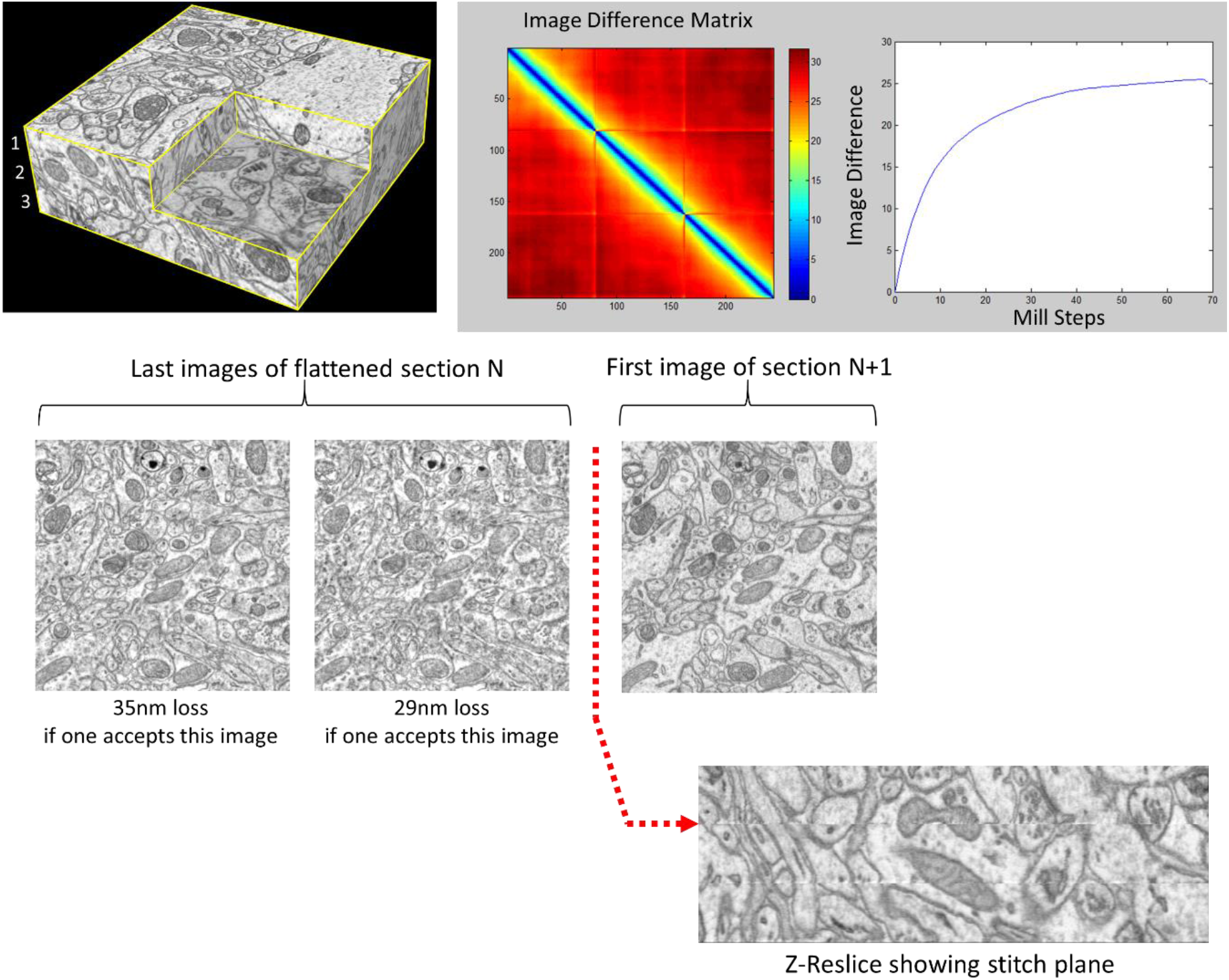
Estimate of sectioning loss for 500nm thick sections of Spurr’s-embedded mouse cortex. (Same dataset shown in **Fig. 2a-c**).

**SOM Figure 23.**
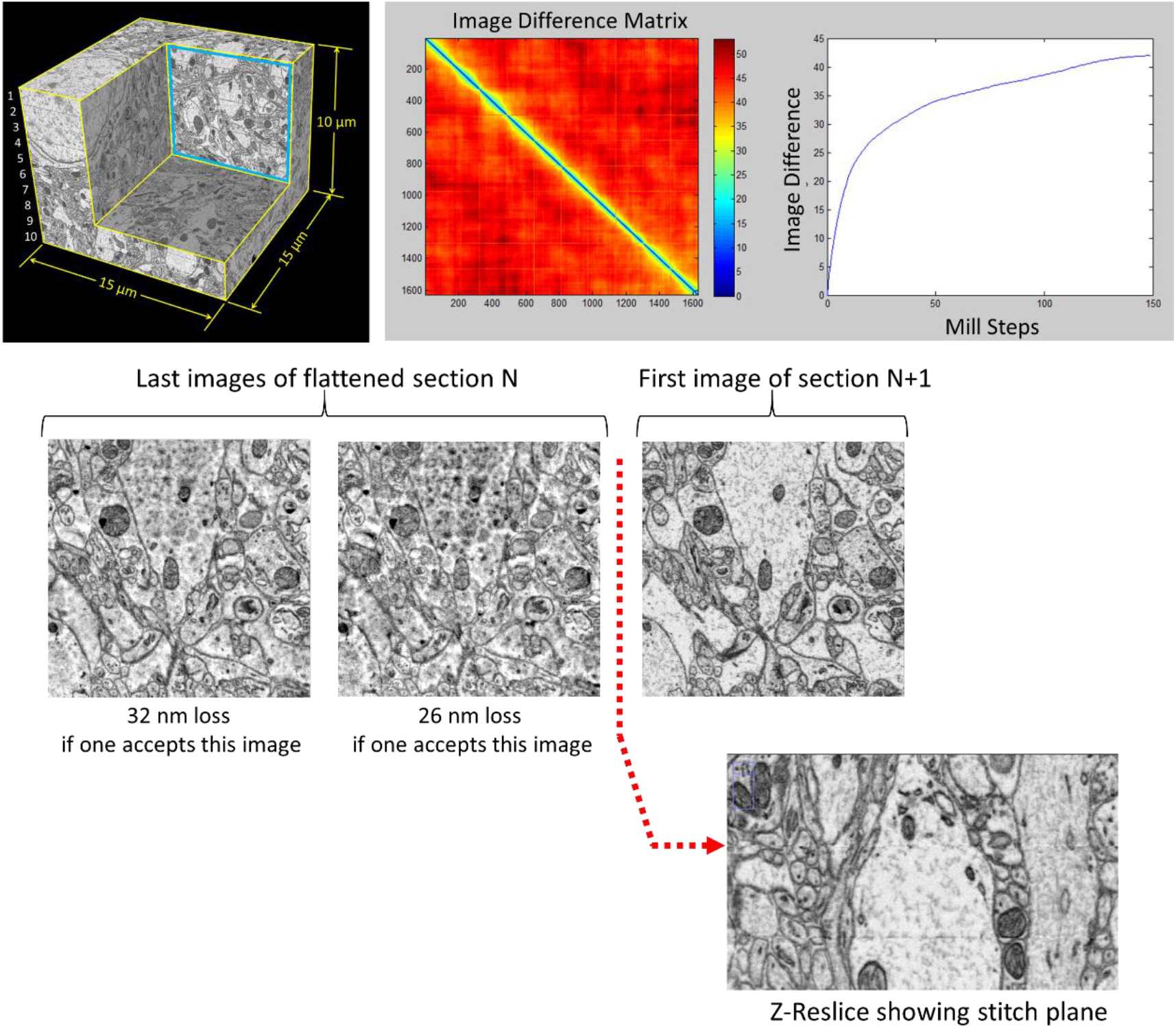
Estimate of sectioning loss for 1 μm thick sections of Spurr’s-embedded mouse cortex tissue (Same dataset shown in **Fig. 2d-f**).

**SOM Figure 24.**
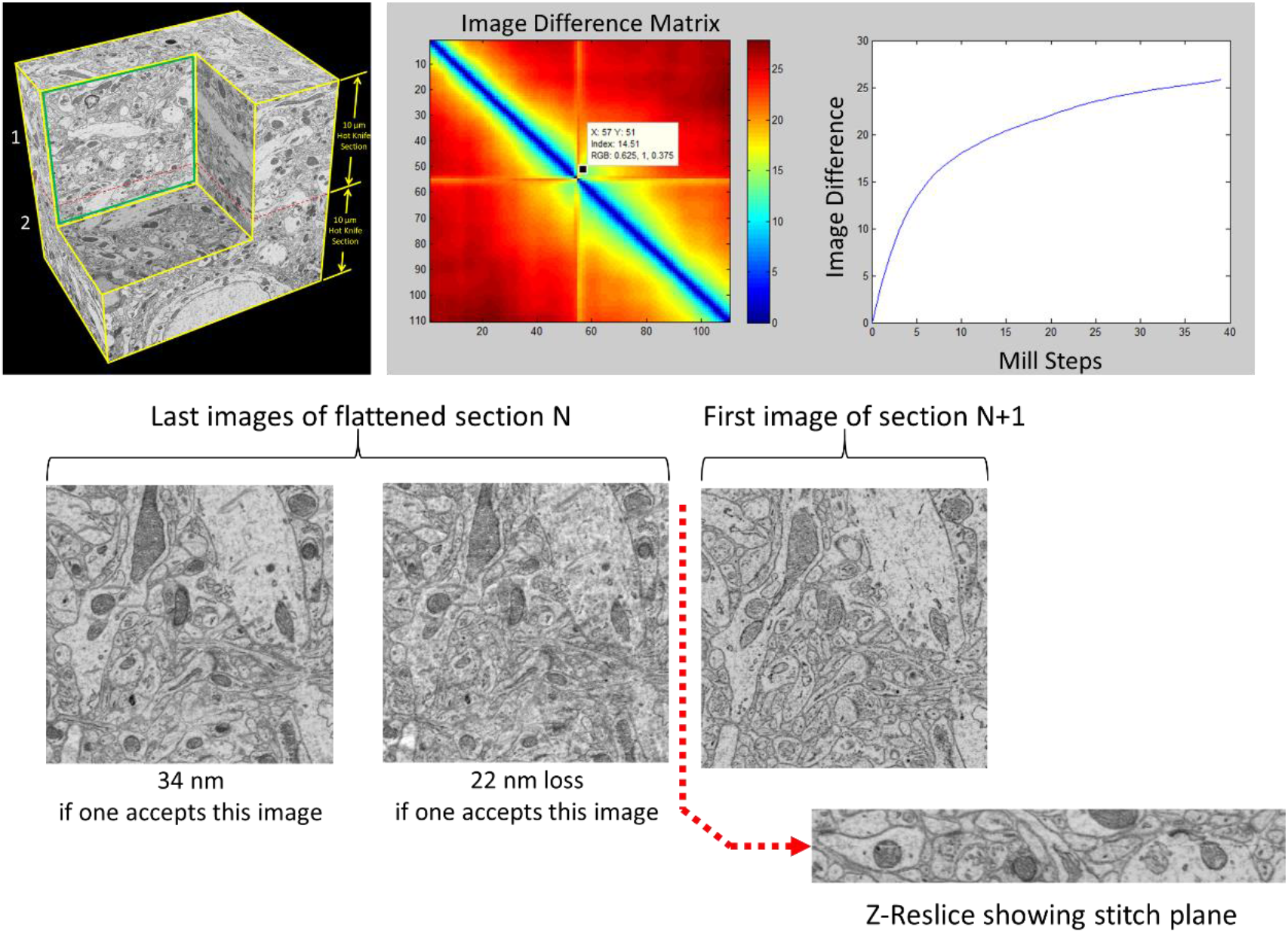
Estimate of sectioning loss for 10 μm thick hot knife sections of Spurr’s-embedded mouse cortex tissue (Same dataset shown in **Fig. 3**).

**SOM Figure 25.**
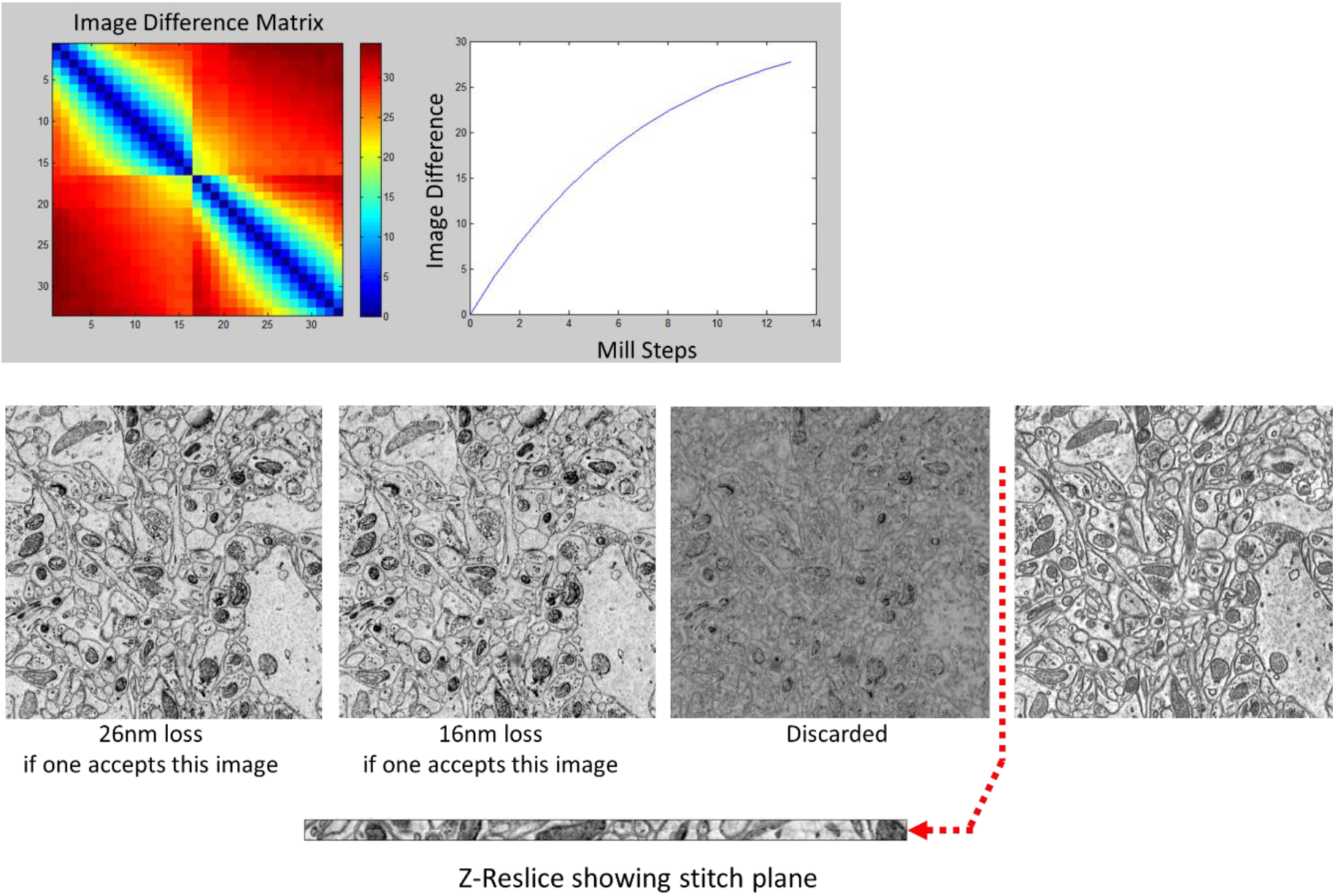
Estimate of sectioning loss for 150 nm thick sections of Spurr’s-embedded mouse cortex tissue.

